# Ultrasound-activated microbubbles mediate F-actin disruptions and endothelial gap formation during sonoporation

**DOI:** 10.1101/2024.08.28.610065

**Authors:** Bram Meijlink, H. Rhodé van der Kooij, Yuchen Wang, Hongchen Li, Stephan Huveneers, Klazina Kooiman

## Abstract

Locally opening up the endothelial barrier in a safe and controlled way is beneficial for drug delivery into the extravascular tissue. Although ultrasound-induced microbubble oscillations can affect endothelial barrier integrity, the mechanism remains unknown. Here we uncover a new role for F-actin in microbubble-mediated endothelial gap formation. Unique simultaneous high-resolution confocal microscopy and ultra-high-speed camera imaging (10 million frames per second) reveal that oscillating microbubbles (radius 1.3-3.8 µm) induce sonoporation in all cells in which F-actin remodeling occurred. F-actin disruption only mainly resulted in tunnel formation (75%) and F-actin stress fiber severing and recoil mainly resulted in cell-cell contact opening within 15 s upon treatment (54%) and tunnel formation (15%). Stress fiber severing occurred when fibers were within reach of the microbubble’s maximum radius during oscillation, requiring normal forces of ≥230 nN. Together, these findings reveal a novel mechanism of microbubble-mediated drug delivery, which associates with the underlying cytoskeletal organization.

## Introduction

The endothelial cells that line the inside of blood vessels form an important physiological barrier between the blood and extravascular tissues, which can hinder the delivery of drugs to diseased extravascular tissue^1, 2^. Having the means to locally open up the endothelial barrier on demand and in a safe and controlled way would be beneficial to get drugs into the extravascular tissue. Lipid-coated microbubbles (diameter of 1-10 µm) with a gas core, originally developed to enhance contrast in diagnostic ultrasound imaging, can be used to locally increase drug delivery^3, 4^. Upon ultrasound insonification, microbubbles oscillate which induces biophysical and mechanical effects on cells^5, 6^. These effects include cellular responses like the perforation of the cell membrane, called sonoporation, which promotes drug uptake into endothelial cells^7, 8^. Oscillating microbubbles can also induce endothelial gaps, either tunnels^9^ through cells or openings between cells^7, 10^, which increases vascular permeability^11, 12^ and thus promote transendothelial drug delivery. Although sonoporation can be predicted by the microbubble behavior^9, 10^, endothelial gap formation cannot^9, 10^. The mechanism by which oscillating microbubbles affect the endothelial barrier integrity therefore remains unclear, which hinders safe and controllable microbubble-mediated transendothelial drug delivery.

Interestingly, recent studies have shown that the filamentous (F)-actin cytoskeleton could be remodeled by oscillating microbubbles, and the resealing of membrane pores induced by sonoporation was synchronized with the recovery of the F-actin cytoskeleton^13, 14^. The actin cytoskeleton is an important regulator of endothelial gap formation and barrier function^15, 16^, suggesting that the F-actin organization could be involved in microbubble-mediated transendothelial drug delivery. Part of the F-actin cytoskeleton consists of contractile actin bundles called stress fibers^17^, which are the result of mechanical stress on a cell^18^. These stress fibers play a crucial role in the formation and rearrangement of cell adhesions^19, 20^ and their contractile forces define cellular shape changes^21^. This makes F-actin stress fibers an interesting cellular component to include in the investigation of the mechanism of microbubble-mediated transendothelial gap formation. So far, the microbubble behavior in relation to F- actin responses has not been investigated^13, 14^ because it requires microsecond time resolution to observe microbubble oscillation in real-time concurrently with actin dynamics.

In this study, we aimed to investigate the role of the F-actin cytoskeleton in transendothelial drug delivery upon oscillating microbubble treatment. The F-actin and cell membrane response to single ultrasound-activated microbubbles was simultaneously visualized at both micrometer and microsecond resolution in live endothelial cells using a unique confocal microscope coupled to an ultra-high-speed camera (10 million frames per second (Mfps)). By examining the responses in the presence and absence of F-actin stress fibers, we found that the severing of F-actin stress fibers by oscillating microbubbles is crucial for creating transendothelial gaps between cells within 15 s upon sonoporation.

## Results

### Oscillating microbubbles induce F-actin disruption and stress fiber severing and recoil

The effect of single oscillating microbubbles with a clinically relevant initial radius, *R0*, ranging from 1.32-3.75 µm on the F-actin and cell membrane was assessed in human umbilical vein endothelial cells (HUVECs) using a custom-built confocal microscope^22^ coupled to an ultra-high-speed camera operated at 10 Mfps (Fig. 1a, b). The F-actin network in the HUVECs was visualized by lentiviral expression of Lifeact-GFP and the cells were cultured to 100% confluency. Propidium iodide (PI) was added to assess sonoporation since PI can only enter viable cells upon membrane disruption and consequently becomes fluorescent due to binding to DNA and RNA^23^. Next, we applied ultrasound and imaged the F-actin cytoskeleton using time-lapse confocal microscopy for 4 min (Fig. 1c). The imaging of 31 different field of views (FOVs) revealed that upon the same magnitude of applied ultrasound (2 MHz, 350 kPa peak negative pressure, 10 cycles), four different F-actin endothelial responses occurred: (a) no disruption, (b) disruption and cytoskeletal recovery, (c) disruption only in the absence of stress fiber severing and (d) stress fiber severing and recoil. For each of these four F-actin responses, a representative example is shown in Fig. 1d-g. A representative example of an oscillating microbubble (*R0* 2.07 µm) that did not disrupt the F-actin nor induce PI uptake is shown in Supplementary Movie 1 and selected frames of this confocal microscopy recording in Fig. 1d. This microbubble was located on a Lifeact-GFP containing cell which was partially attached to one of its non-Lifeact-GFP containing neighboring cells before ultrasound application as observed from the cell membrane staining (indicated with yellow arrow). Upon ultrasound, cell-cell contact opening occurred ∼40 s after treatment at the location of the partial attachment to the neighboring cell (indicated with yellow arrowheads). The microbubble oscillation amplitude, defined as the microbubble’s maximum radius during oscillation (*Rmax*) minus the microbubble’s initial radius (*R0*), was 0.88 µm. Z-stack confocal microscopy after the time-lapse imaging revealed the absence of sonoporation and F-actin disruption (Supplementary Fig. 1a). Figure 1e shows the representative example of immediate disruption of the F-actin (i.e., reduction in green fluorescent signal) by the oscillating microbubble (*R0* 1.96 µm) with a 1.43 µm oscillation amplitude at the location of the microbubble and PI uptake (i.e., sonoporation). The disrupted F-actin cytoskeleton recovered within the time-lapse confocal microscopy recording (Supplementary Movie 2) and PI uptake stabilized after 6 s (Fig. 1e), indicating a resealed cell membrane pore as previously reported^10, 24^. The resealed membrane pore was confirmed by z-stack confocal microscopy (Supplementary Fig. 1b). As shown in Fig. 1f and Supplementary Movie 3, the F-actin could also disrupt and remain disrupted over time at the location of the microbubble and PI uptake. This microbubble had an *R0* of 2.15 µm and oscillation amplitude of 2.47 µm. Forty-two s after treatment, PI uptake stabilized, indicating a resealed cell membrane pore, but the cell membrane signal did not recover at the location of the microbubble. Z-stack confocal microscopy revealed a transendothelial tunnel had formed (Supplementary Fig. 1c) with intact cell membranes along the tunnel. In Fig. 1g, unlike in the other representative examples shown in Fig. 1d-f, the oscillating microbubble (*R0* 1.68 µm) with a 1.96 µm oscillation amplitude was located on the F-actin stress fibers, resulting in direct severing of the fibers closest to microbubble location. Upon severing, the stress fiber recoiled along the longitudinal direction towards the end of the initial stress fibers. Upon stress fiber severing and recoil, a strong retraction of the cell was observed within 9 s of the ultrasound treatment leading to cell-cell contact opening (Supplementary Movie 4). PI uptake in this cell was also observed. Z-stack confocal microscopy after the time-lapse imaging showed the creation of a transendothelial gap where the cell retracted, only leaving behind some of the membrane (Supplementary Fig. 1d). Taken together, these experiments show that there is a major difference in the endothelial cell response to ultrasound-activated microbubbles, which associates with the underlying cytoskeletal organization of the stress fibers.

**Figure 1.**
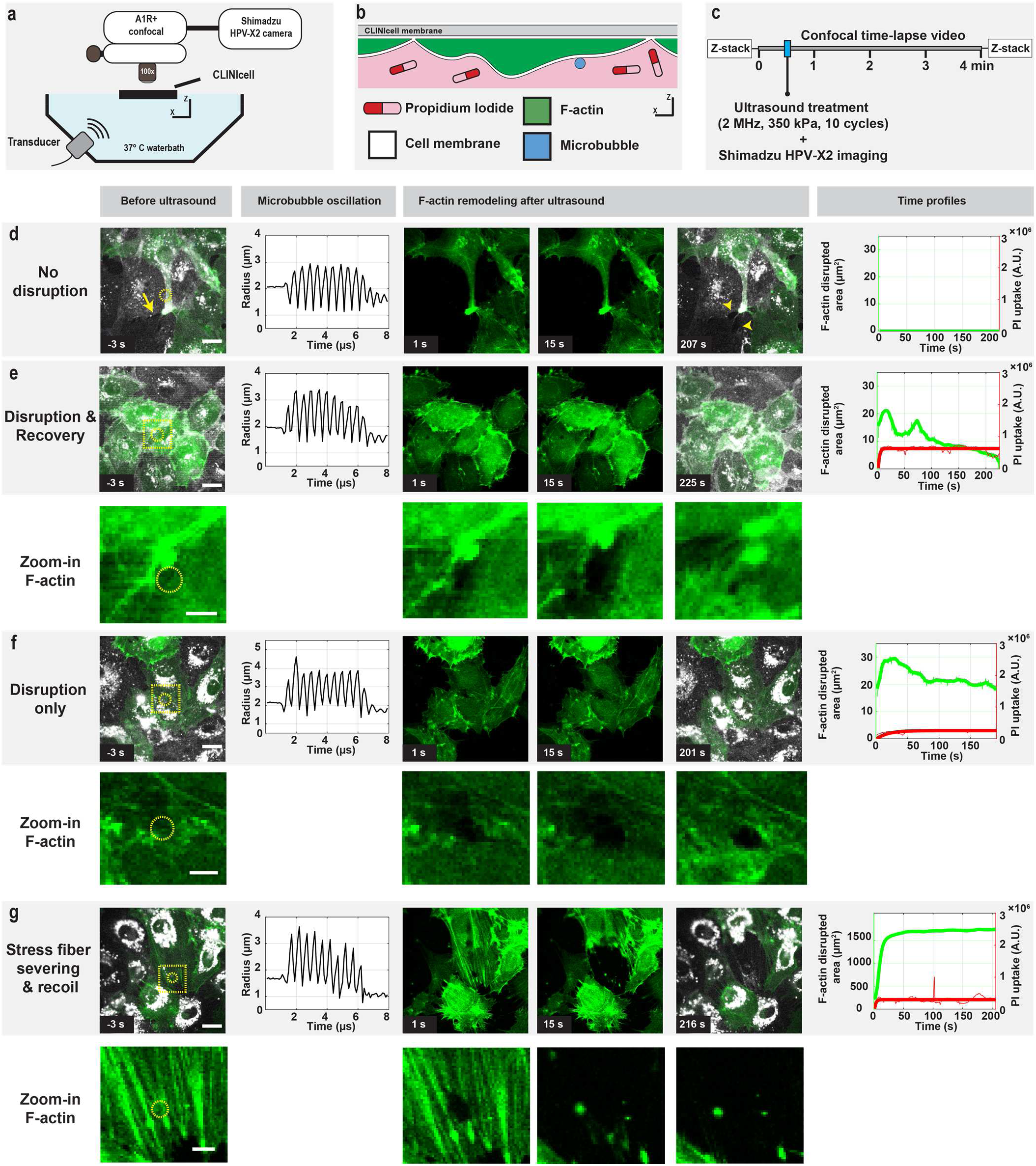
| Experimental set-up and F-actin remodeling upon oscillating microbubble treatment. (**a**) Experimental set-up (not drawn to scale): confocal microscope coupled to the Shimadzu HPV-X2 ultra-high-speed camera (10 Mfps) used to visualize the cellular and microbubble responses. The transducer in the water bath generated the ultrasound (**b**) Schematic representation of the fluorescently labeled F-actin (green) and cell membrane (white) in the HUVECs, microbubble (blue), and model drug propidium iodide (red) within the CLINIcell in the orientation used during the experiment. (**c**) Imaging timeline of the experiment. (**d-g**) Representative images of the observed types of F-actin remodeling with zoom-in of the F-actin when remodeling was observed. For every example, the images from left to right show: a confocal image of the F-actin (pseudo-colored in green) and cell membrane (pseudo-colored in white) before ultrasound with a dotted yellow circle indicating the microbubble location and yellow dotted square indicating the zoom-in location of the F-actin remodeling; the microbubble radius-time curve obtained from analysis of the ultra-high-speed-imaging during microbubble oscillation; two additional confocal images of the F-actin remodeling at different timepoints after treatment; one additional confocal image of the F- actin remodeling and cell membrane at the end of the 4-min time-lapse; and quantification of the time profiles of the disrupted F-actin area (green, thick line represents moving average) and propidium iodide (PI) fluorescent intensity (red, thick line shows previously reported fit^9, 10^) in the cell, where the time is represented compared to the ultrasound insonification. Scale bars represent 20 µm, except for the zoom-ins where it represents 5 µm. The 3D confocal microscopy images of these examples before and after time-lapse confocal microscopy imaging are shown in Supplementary Fig. 1. (**d**) No F-actin remodeling and no sonoporation (corresponding confocal microscopy recording in Supplementary Movie 1). Yellow arrow indicates where the Lifeact-GFP-containing cell is partially attached to one of its non-Lifeact-GFP-containing neighboring cells. Yellow arrowheads indicate the location of cell-cell contact opening. (**e**) F-actin disruption followed by cytoskeletal recovery (i.e., disappearance & recovery of the green signal) and sonoporation (i.e., PI uptake) (corresponding confocal microscopy recording in Supplementary Movie 2). (**f**) F-actin disruption only that remained present, sonoporation and formation of a transendothelial tunnel (corresponding confocal microscopy recording in Supplementary Movie 3). (**g**) F-actin stress fiber severing and recoil, sonoporation and cell-cell contact opening (corresponding confocal microscopy recording in Supplementary Movie 4). Note that the time profile for the F-actin disrupted area (i.e., left y-axis) has a larger range than the time profiles in d-f.

### F-actin remodeling by oscillating microbubble induces intracellular and transendothelial drug delivery

Quantification of the F-actin disrupted area revealed that the initially disrupted area ∼1 s after ultrasound application (Fig. 2a) and maximally disrupted area (Fig. 2b) were significantly larger for the F- actin remodeling categories disruption only and stress fiber severing and recoil than for the category of F-actin disruption and recovery. No significant differences were detected for the initially and maximally disrupted F-actin areas between the remodeling categories disruption only and stress fiber severing and recoil. The median maximally disrupted F-actin area was 2.4 times as large as the median initially disrupted F-actin area for the cells in which the oscillating microbubble induced F-actin disruption and recovery, while this was 1.4 times as large for disruption only and 7.0 times as large for the stress fiber severing and recoil categories. In addition, the median (with interquartile range (IQR) between brackets) maximally disrupted F-actin area of 22 (8-29) µm^2^ for the F-actin remodeling category of disruption and recovery, was similar to the initially disrupted F-actin area for the F-actin remodeling categories of disruption only (28 (21-37) µm^2^) and stress fiber severing and recoil (27 (14-46) µm^2^). To link F-actin remodeling upon oscillating microbubbles with drug delivery outcome, we quantified both distinct types of drug delivery routes: intracellular and transendothelial drug delivery. Fig. 2c shows that membrane pore formation, i.e., sonoporation, was induced in all cells in which F-actin remodeling occurred. In 92% (23 out of 25) of these cells, the induced membrane pore resealed, while in the remaining cells the induced membrane pore did not reseal. The cells with non-resealing membrane pores were all in the F- actin remodeling category of stress fiber severing and recoil.

**Figure 2.**
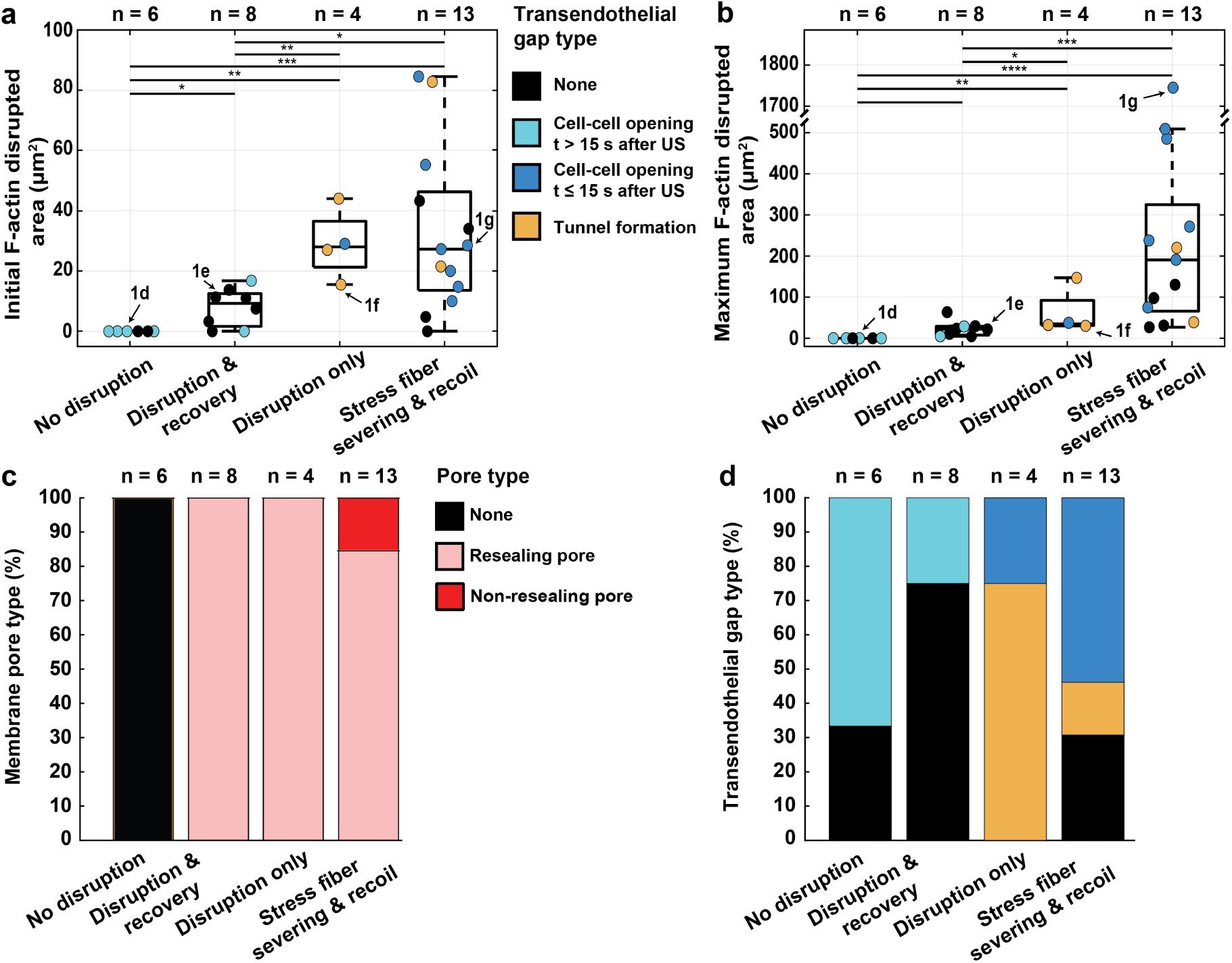
| Cellular and drug delivery responses upon F-actin disruption by oscillating microbubbles. (**a**) Quantification of initial disrupted F-actin area at ∼1 s after ultrasound treatment. (**b**) Quantification of maximum disrupted F-actin area observed. Note that the y-axis has a larger range than in (a) and is discontinued into two sections of 0-500 and 1700-1800 µm^2^. (**c**) Quantification of induced resealing or non-resealing cell membrane pore by oscillating microbubbles per type of F-actin remodeling. (**d**) Quantification of induced transendothelial gap types tunnel and cell-cell contact opening by oscillating microbubbles per type of F-actin remodeling. (**a-b**) Significance is indicated with * (P < 0.05), ** (P < 0.01), *** (P < 0.001) and **** (P<0.0001). Boxplots represent the median and the boxes indicate the 25th and 75th percentiles with whiskers ranging from minimum to maximum values excluding outliers. Datapoints which correspond to the examples shown in Fig. 1d-g are indicated with arrows. US = Ultrasound. (**a-d**) N-numbers represent individually investigated FOVs.

As shown in Fig. 2d, transendothelial gap formation upon F-actin remodeling could occur in two ways, namely by a tunnel (i.e., transcellular perforation within one cell; indicated in orange) or opening between cells (i.e., cell-cell junction opening; indicated in light and dark blue). Tunnel formation was only observed in cells in which the F-actin was disrupted only (3 out of 4 cells; 75% occurrence) and in cells in which stress fiber severing and recoil occurred (2 out of 13 cells; 15% occurrence). Cell-cell contact (junction) opening occurred across all F-actin remodeling categories, including when the F-actin was not disrupted (Fig. 2d). Six out of the 31 ultrasound and microbubble treated cells (19%) already had a gap between them and their neighboring cells before the application of ultrasound. The number of cells which already had a gap before control treatment, was four out of 14 (29%) in the ultrasound only and five out of 21 (24%) for the microbubble only controls. In cells with existing gaps before ultrasound and microbubble treatment, the opening of cell-cell contacts occurred significantly later than the <15 s for cells that were fully attached to their neighbors (Supplementary Fig. 2a). For this reason, cell-cell contact opening was classified as slow opening (i.e., > 15 s after ultrasound; light blue in Fig. 2d) or fast opening (i.e., ≤ 15 s after ultrasound; dark blue in Fig. 2d). The slow cell-cell contact opening occurred in 6 out of the 31 studied cells (19%) and was induced in cells in which F-actin was not disrupted (4 out of 6 cells; 67%) or was disrupted and recovered upon ultrasound and microbubble treatment (2 out of 8 cells; 25%), as well as in 24% of the cells in which F-actin was not disrupted in the control microbubble only (5 out of 21 cells) and ultrasound only (3 out of 14 cells; 21%) treatments. The location of the slow opening of the cell-cell junctions was not in the vicinity of the microbubble. The fast cell-cell contact opening was induced in cells in which the F-actin was disrupted only (1 out of 4 cells for microbubble located in preexisting gap between cells (Supplementary Movie 5); 25% occurrence) or F-actin stress fiber severing and recoil occurred (7 out of 13 cells; 54% occurrence). All but one of the cells in which fast cell-cell contact opening occurred were fully attached to their neighbors before the ultrasound and microbubble treatment. Thus, oscillating microbubbles induce sonoporation in all cells in which F-actin remodeling occurred, where F-actin disruption only mainly resulted in tunnel formation (75%) and F-actin stress fiber severing and recoil mainly resulted in cell-cell contact opening (54%) and tunnel formation (15%).

### Oscillating Microbubble behavior

The oscillating microbubbles that induced the F-actin remodeling responses were located up to 9.5 µm from the cell edge (Fig. 3a). The microbubbles that induced F-actin stress fiber severing and recoil were not positioned significantly closer or farther from the cell edge than the microbubbles that induced the other F-actin responses (Fig. 3a), suggesting stress fiber severing cannot be predicted based on the initial microbubble location on the cell. There was no significant difference in the initial radius of the microbubble between the F-actin remodeling responses (Supplementary Fig. 2b). As shown in Fig. 3b, the analysis of the microbubble oscillation showed a significantly lower microbubble oscillation amplitude when the F-actin was not disrupted in comparison to the three F-actin remodeling responses. When the F-actin was not disrupted by the oscillating microbubble, the microbubble oscillation amplitude was smaller than the previously reported sonoporation threshold of 0.9 µm^9^. The three F-actin remodeling responses were induced with microbubble oscillation amplitudes > 0.9 µm. However, no significant differences in microbubble oscillation amplitude were observed between the three F-actin remodeling responses. Quantification of the asymmetry of the microbubble oscillation, defined as the ratio of microbubble expansion over compression (*E*/*C*)^10^, showed that the microbubbles oscillated symmetrically (0.5 ≤ *E*/*C* ≥ 2) and with expansion only (*E*/*C* > 2). The microbubbles that induced the three types of F-actin remodeling showed significantly higher expansion during oscillation compared to the non-F-actin-remodeling microbubbles (Supplementary Fig. 2c). As shown in Fig. 3c, the majority of the microbubbles fragmented during oscillation when the F-actin was remodeled, namely in 63% (5 out of 8) of the cells with F-actin disruption and recovery, in 100% of the cells with F-actin disruption only and 85% (11 out of 13) of the cells with F-actin stress fiber severing and recoil. An example of a microbubble that fragmented during oscillation is shown in Supplementary movie 6 with selected frames of the ultra-high-speed recording presented in Fig. 3d. This microbubble induced F-actin stress fiber severing and recoil as shown in Fig. 1g. In addition, two of the oscillating microbubbles showed 3^rd^ order shape modes, meaning a triangular shape during oscillation, as illustrated in Fig. 3e and Supplementary Movie 7. One of these microbubbles did not induce F-actin disruption, while the other induced F-actin disruption and recovery. In conclusion, the microbubble oscillation amplitude needs to be > 0.9 µm to induce sonoporation and F-actin remodeling, but there is no correlation between the microbubble oscillation amplitude and type of F- actin remodeling. Also, the oscillating microbubbles do not need to be near the cell edge to induce transendothelial gaps between cells.

**Figure 3.**
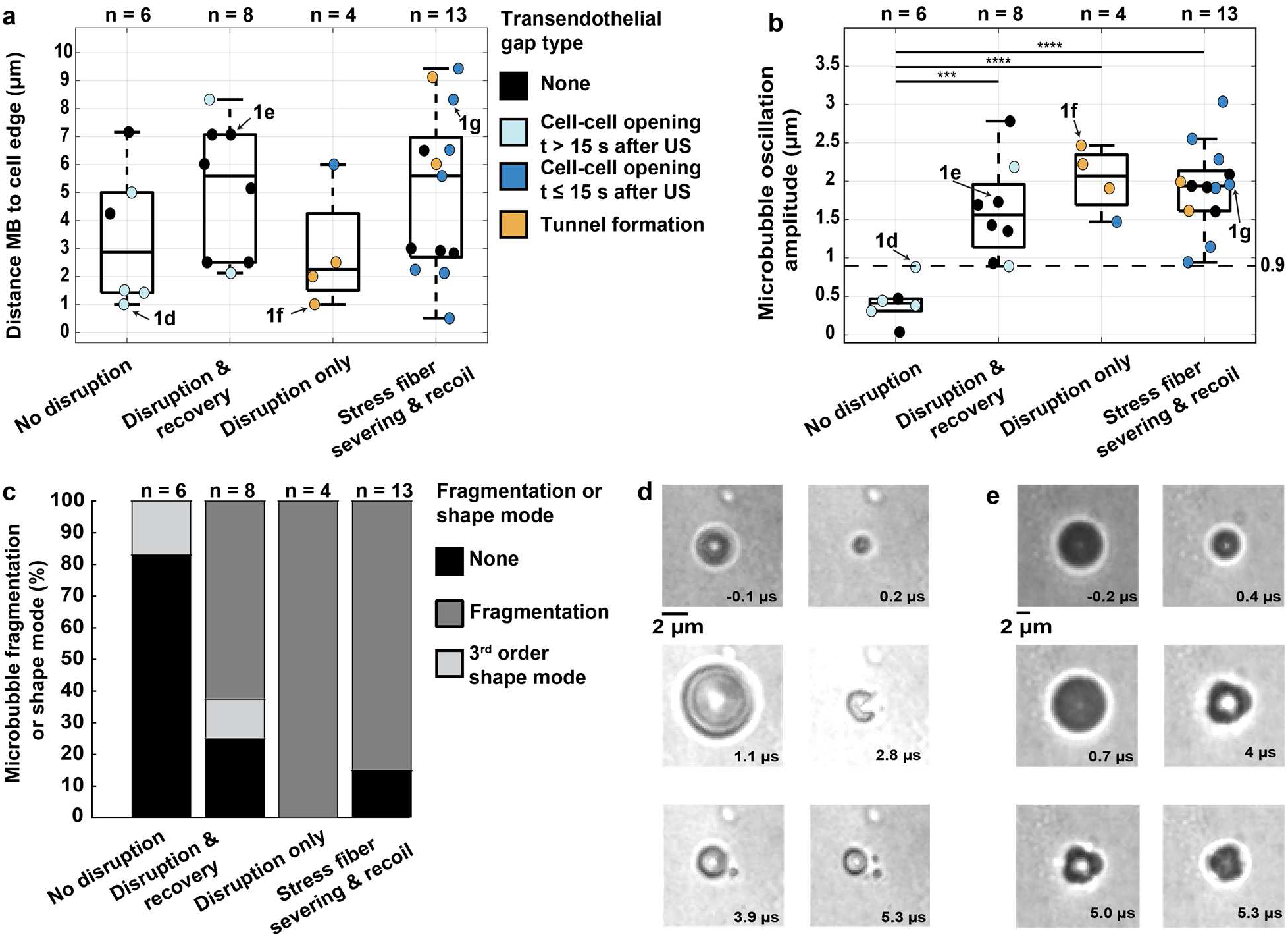
| Microbubble oscillation behavior. (**a**) Quantification of the distance between the center of the microbubble (MB) and cell edge before ultrasound application per type of F-actin remodeling. (**b**) Quantification of the microbubble oscillation amplitude (*R_max_* - *R_0_*) per type of F-actin remodeling where dotted line at 0.9 µm indicates the sonoporation threshold. (**a, b**) Boxplots represent the median and the boxes indicate the 25th and 75th percentiles with whiskers ranging from minimum to maximum values excluding outliers. Significance is indicated with *** (P < 0.001) and **** (P<0.0001). Datapoints which correspond to the examples shown in Fig. 1d-g are indicated with arrows. US = Ultrasound. (**c**) Quantification of fragmentation and 3^rd^ order shape modes during microbubble oscillation per type of F-actin remodeling. (**a-c**) N-numbers represent individually investigated FOVs. (**d**) Selected frames of the ultra-high-speed camera recording of a microbubble that fragmented during oscillation, corresponding to the example shown in Fig. 1g where the F-actin stress fibers were severed and recoiled. See Supplementary Movie 6 for the ultra-high-speed recording. (**e**) Selected frames of an ultra-high-speed camera recording of a microbubble that had 3^rd^ order shape modes during oscillation, corresponding to the example shown in Fig. 4e where the F-actin disrupted and recovered. See Supplementary Movie 7 for the ultra- high-speed recording. (**e-f**) Time is indicated compared to the start of the ultrasound application. Scale bars represent 2 µm.

### F-actin stress fibers severing occurs in close proximity to oscillating microbubbles

We set out to investigate why oscillating microbubbles-induced stress fiber severing and recoil in some cells while not in others. Assessment of the number of pre-existing stress fibers per cell (Fig. 4a) and cell area (Supplementary Fig. 3a) revealed no significant differences between no F-actin disruption and the three different F-actin responses induced by oscillating microbubbles. To analyze the cytoskeletal organization of F-actin stress fibers around the location of the microbubbles, we measured the distance from the microbubble center to the nearby pre-existing stress fibers before applying ultrasound using a spherical coordinate system centered at the microbubble, with segmentations into radial bands at 3 µm intervals (Fig. 4b). Fig. 4c shows that the stress fibers that were severed (indicated in green) were within 4.7 µm from the microbubble center, which was significantly closer in comparison to the stress fibers that were not severed (indicated in black). Although this indicates that the proximity of a microbubble to a stress fiber is a contributing factor, it is not the sole determinant as 12 out of the 31 non-served fibers were also within 4.7 µm from the microbubble center. As stress fiber severing occurred beyond the initial microbubble radius (dotted line in Supplementary Fig. 3b), we assessed if there was a relation between stress fiber severing and the microbubble’s maximum radius during oscillation. Of the severed stress fibers, 89% (25 out of 28) were within the microbubble’s maximum radius during oscillation (located below the dotted line in Fig. 4d), while 11% (3 out of 28) were even above the microbubble’s maximum radius (i.e., located above the dotted line in Fig. 4d). Of the non-severed stress fibers, only 2.7% (7 out of 259) were within the microbubble’s maximum radius (located below the dotted line in Fig. 4d). Two of these non-severed stress fibers, located within the same cell, are indicated by black diamond-shaped symbols below the dotted line in Fig. 4d. This microbubble (*R0* 3.8 µm) exhibited 3^rd^ order shape modes as shown in Fig. 3e (Supplementary movie 7). Interestingly, this oscillating microbubble was able to induce sonoporation and F-actin disruption in between two stress fibers followed by recovery, as shown in the selected fluorescent images and time profile quantification in Fig. 4e (Supplementary movie 8). Z-stack confocal microscopy confirmed recovery of the F-actin without stress fiber severing (Supplementary Fig. 4). Calculations of the generated normal forces by the oscillating microbubbles are presented in Fig. 4f. The normal forces were between 230 and 8,460 nN with a median of 1,500 nN and interquartile range of 390-2,300 nN. Together, our data show that oscillating microbubbles sever F-actin stress fibers when they are in reach of the microbubble’s maximum radius during oscillation and generate normal forces of ≥230 nN.

**Figure 4.**
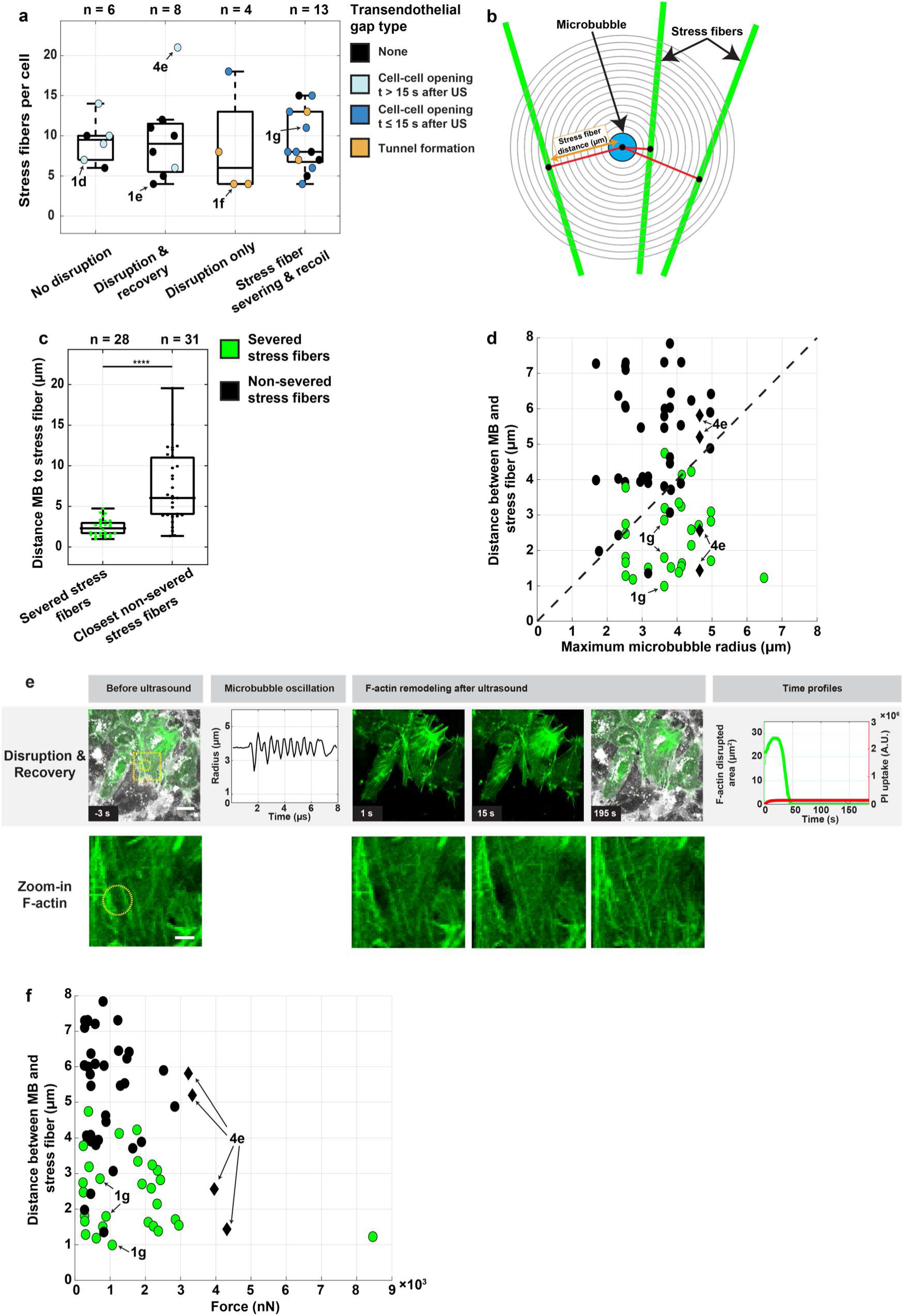
| F-actin stress fiber severing by oscillating microbubbles. (**a**) Quantification of the number of F-actin stress fibers in each cell of interest per type of F-actin remodeling. (**b**) Schematic representation of polar coordinate pattern used to quantify the distance between the microbubble center and all stress fibers in the cell of interest. (**c**) Quantification of the distances between the microbubble center to all severed and closest non-served stress fibers in the cell of interest. (**d**) Relation for distance between the microbubble (MB) center and surrounding stress fibers and maximum microbubble radius (*R_max_*) during oscillation. Since both parameters are quantified from the center of the MB, a theoretical line (black dotted) can be drawn that separates the F-actin stress fibers that were within reach of the *R_max_* (below dotted line) from the stress fibers that were outside the *R_max_* reach (above dotted line). The black diamond data points indicate microbubbles that showed 3^rd^ order shape modes during oscillation. (**e**) Representative example of F-actin disruption and recovery by a microbubble that showed 3^rd^ order shape mode during oscillation (diamond-shaped data points in Fig. 4d). The images from left to right show: a confocal image of the F-actin (pseudo-colored in green) and cell membrane (pseudo-colored in white) before ultrasound with a dotted yellow circle indicating the microbubble location and yellow dotted square indicating the zoom-in location of the F-actin remodeling; the microbubble radius-time curve obtained from analysis of the ultra-high-speed- imaging (see Supplementary Movie 7 and Fig. 3e for selected frames of this recording) during microbubble oscillation; two additional confocal images of the F-actin remodeling at different timepoints after treatment (see Supplementary Movie 8); one additional confocal image of the F-actin remodeling and cell membrane at the end of the 4-min time-lapse; and quantification of the time profiles of the disrupted F-actin area (green, thick line represents moving average) and propidium iodide (PI) fluorescent intensity (pseudo-colored in red, thick line shows previously reported fit^9, 10^) in the cell, where the time is represented compared to the ultrasound insonification. Scale bars represent 20 µm, except for the zoom-ins where it represents 5 µm. See Supplementary Fig. 4 for the 3D z-stack confocal microscopy imaging before and after time-lapse confocal microscopy imaging. (**f**) Relation for the distance between the MB center and surrounding stress fibers and the generated normal force by the oscillating microbubbles on the stress fibers. (**a, c**) N-numbers represent individually investigated FOVs. Boxplots represent the median and the boxes indicate the 25th and 75th percentiles with whiskers ranging from minimum to maximum values excluding outliers. Significance is indicated with **** (P<0.0001). (**a, d, f**) Datapoints which correspond to the examples shown in Fig. 1d-g and 4e are indicated with arrows.

### Oscillating microbubble-induced cell-cell contact opening does not occur in the absence of F-actin stress fibers

To investigate whether the presence of F-actin stress fibers was required to induce cell-cell contact opening upon oscillating microbubble-induced sonoporation, cells were incubated with the ROCK inhibitor Y-27632 for 1 h prior to imaging to remove stress fibers. Upon microbubble and ultrasound treatment of 17 different FOVs, three different F-actin endothelial responses occurred: (a) no disruption, (b) disruption and recovery, and (c) disruption only. Figure 5a shows a representative example of F-actin disruption and recovery by an oscillating microbubble (*R0* 1.85 µm) with a 1.94 µm oscillation amplitude at the location of the microbubble and PI uptake (i.e., sonoporation). The disrupted F-actin cytoskeleton recovered within the time-lapse confocal microscopy recording and PI uptake stabilized after 5 s, indicating a resealed cell membrane pore (Supplementary Movie 9). The resealed membrane pore was confirmed by z-stack confocal microscopy (Supplementary Fig. 5a). The oscillating microbubble (*R0* 1.77 µm) with a 1.13 µm oscillation amplitude that induced sonoporation and F-actin disruption only is presented in Supplementary Movie 10 and selected frames of this confocal microscopy recording are shown in Fig. 5b. Eighteen s after ultrasound application, PI uptake stabilized, indicating a resealed cell membrane pore. However, the F-actin and cell membrane signal did not recover at the location of the microbubble. Z-stack confocal microscopy confirmed the formation of a transendothelial tunnel (Supplementary Fig. 5b).

**Figure 5.**
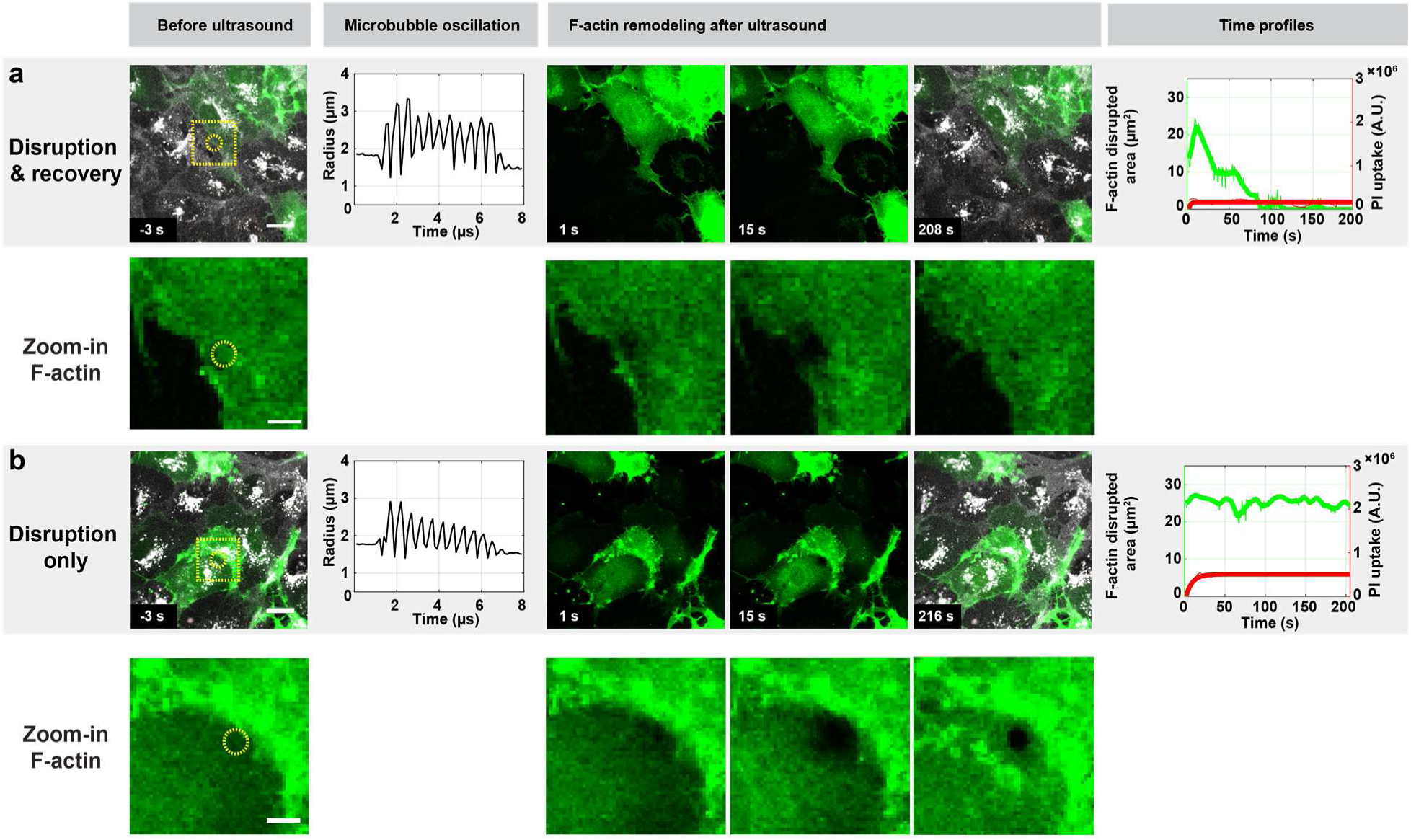
| F-actin remodeling upon oscillating microbubble treatment in Y-27632 incubated cells which removes F-actin stress fibers. (**a-b**) The representative images from left to right show: a confocal image of the F-actin (pseudo-colored in green) and cell membrane (pseudo-colored in white) before ultrasound with a dotted yellow circle indicating the microbubble location and yellow dotted square indicating the zoom-in location of the F-actin remodeling; the microbubble radius-time curve obtained from analysis of the ultra-high-speed-imaging during microbubble oscillation; two additional confocal images of the F-actin remodeling at different timepoints after treatment; one additional confocal image of the F-actin remodeling and cell membrane at the end of the 4-min time-lapse; and quantification of the time profiles of the disrupted F-actin area (green, thick line represents moving average) and propidium iodide (PI) fluorescent intensity (red, thick line shows previously reported fit^9, 10^) in the cell, where the time is represented compared to the ultrasound insonification. Scale bars represent 20 µm, except for the zoom-ins where it represents 5 µm. The 3D confocal microscopy images of these examples before and after time-lapse confocal microscopy imaging are shown in Supplementary Fig. 5. (**a**) F-actin disruption and recovery and sonoporation (corresponding confocal microscopy recording in Supplementary Movie 9) (**b**) F-actin disruption only that remained present, sonoporation and formation of a transendothelial tunnel (corresponding confocal microscopy recording in Supplementary Movie 10).

For Y-27632-incubated cells, quantification of the F-actin disrupted area showed no significant differences for both the initial (i.e., ∼1 s after ultrasound, Fig. 6a) and maximum (Fig. 6b) disrupted areas between no F-actin disruption and the F-actin remodeling categories. The median maximally disrupted F- actin area was 3.25 times larger than the median initially disrupted F-actin area for the cells in which the oscillating microbubble induced F-actin disruption and recovery, while this was 2.97 times larger for the disruption only category. When comparing cells without and with Y-27632 incubation, the median (with interquartile range (IQR) between brackets) initial F-actin disrupted area was significantly higher for cells without (28 (21-37) µm^2^; Fig. 2a) than cells with Y-27632 incubation (5 (1-13) µm^2^) for the F-actin remodeling category of disruption only. Both distinct types of drug delivery routes, i.e., intracellular and transendothelial drug delivery, were quantified and linked with the F-actin remodeling upon oscillating microbubbles. Similarly to the non-Y-27632 incubated cells, membrane pore formation, i.e., sonoporation, was induced in all Y-27632 incubated cells in which F-actin remodeling occurred (Fig. 6c). In addition, all induced membrane pores resealed. As shown in Fig. 6d, transendothelial gap formation occurred in only one cell in which the F-actin was disruption only (1 out of 6 cells; 17% occurrence, example in Fig. 5b and z-stack confocal microscopy in Supplementary Fig. 5b) in the form of a tunnel. Interestingly, in the Y-27632-incubated cells no fast and slow cell-cell contact opening were observed during oscillating microbubble treatment as well as the microbubble only control treatment, even though existing gaps between cells were present before treatment (n=1 in ultrasound plus microbubble and n=2 in microbubble only treatment groups).

**Figure 6.**
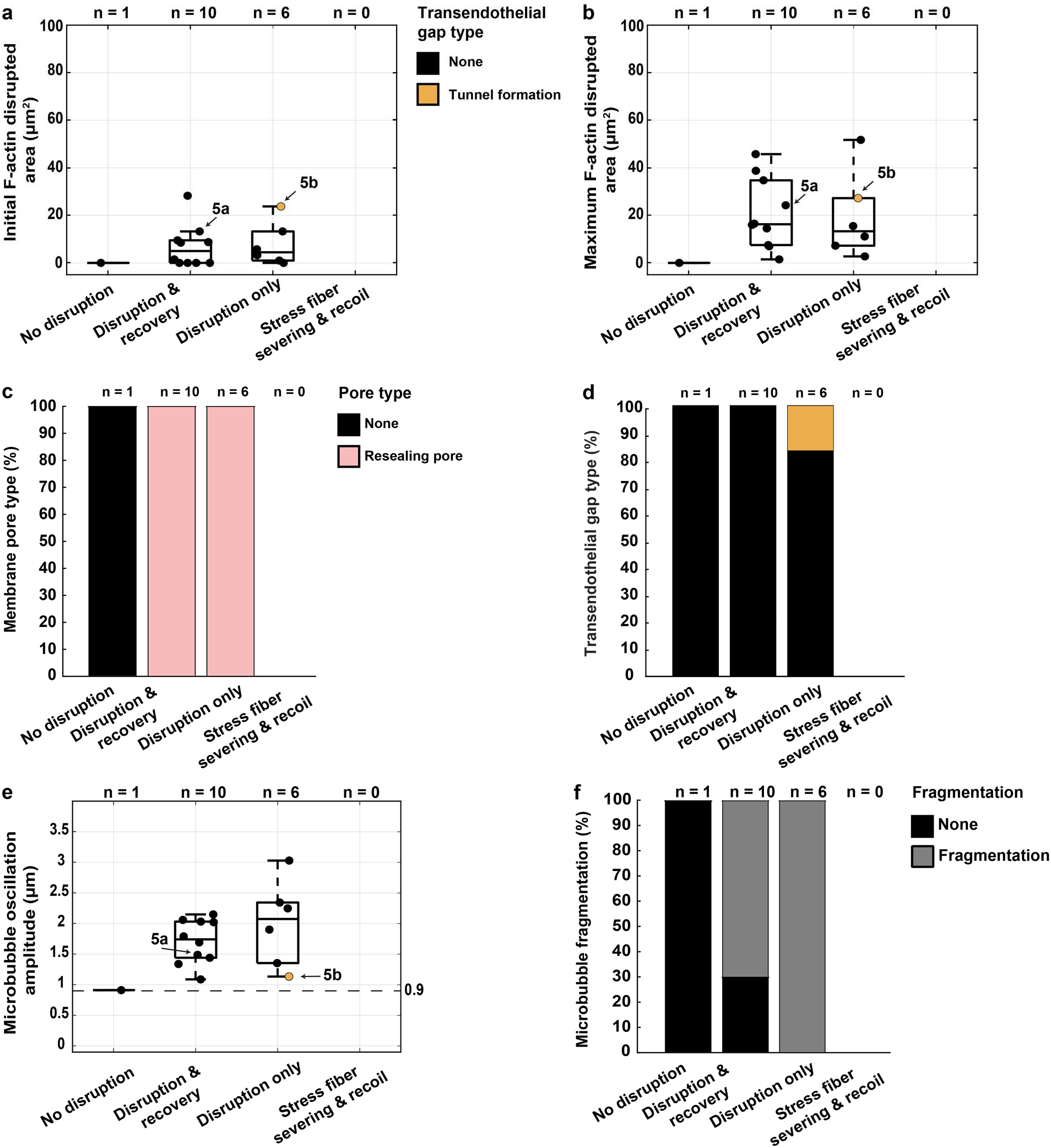
| Cellular and drug delivery responses upon F-actin disruption by oscillating microbubble in Y-27632 incubated cells which removes F-actin stress fibers. (**a**) Quantification of initial disrupted F-actin disrupted area at ∼1 s after ultrasound treatment. (**b**) Quantification of maximum F-actin disrupted area observed. (**c**) Quantification of induced resealing or non-resealing cell membrane pore by oscillating microbubble per type of F-actin remodeling. (**d**) Quantification of induced transendothelial gap type tunnel by oscillating microbubble per type of F- actin remodeling. (**e**) Quantification of the microbubble oscillation amplitude (*R_ma_*_x_ - *R_0_*) during oscillation per type of F-actin remodeling. (**f**) Quantification of fragmentation during microbubble oscillation per type of F-actin remodeling. (**a**, **b**, **e**) No significant differences were observed. Boxplots represent the median and the boxes indicate the 25th and 75th percentiles with whiskers ranging from minimum to maximum values excluding outliers. Datapoints which correspond to the examples shown in Fig. 5a-b are indicated with arrows. (**a-f**) N-numbers represent individually investigated FOVs.

The microbubble’s initial radius, oscillation amplitude (Supplementary Fig. 6) and *E*/*C* (Supplementary Fig. 7a) were similar for cells without and with Y-27632 incubation. In contrast to the cells without Y- 27632 incubation, we found significant differences between the initial radius of the microbubble that did not induce F-actin disruption (*R0* 3 µm) compared to the microbubbles that induced F-actin disruption and recovery (median and IQR radius of 1.8 (1.5-2.0) µm) and F-actin disruption only (1.9 (1.8-2.5) µm) in the Y-27632-incubated cells (Supplementary Fig. 7b). However, there were no significant differences for the microbubble oscillation amplitudes between the F-actin remodeling categories (Fig. 6e). All microbubbles that induced F-actin remodeling and sonoporation had an oscillation amplitude larger than the sonoporation threshold of 0.9 µm, while the oscillation amplitude of the microbubble that did not induce F-actin disruption was equal to 0.9 µm. Similar to the non-Y27632-incubated cells, the majority of the microbubbles in the Y-27632-incubated cells fragmented during oscillation when the F-actin was remodeled, namely in 70% (7 out of 10) of the F-actin disruption and recovery and 100% of the F-actin disruption only categories (Fig. 6f). In conclusion, after the removal of F-actin stress fibers using Y-27632- incubation, sonoporation and F-actin disruption could still be induced and the microbubble oscillations were not affected. However, transendothelial tunnel formation was reduced and cell-cell contact opening was completely absent.

## Discussion

Several recent studies show the importance of oscillating microbubbles for local drug delivery as shown by Ozdas et al.^25^ and in recent reviews^26, 27^. The response of an endothelial cell to an oscillating microbubble is pivotal for safe and controlled drug delivery. Although the microbubble oscillation amplitude is a predictor for sonoporation, it does not predict the formation of endothelial gaps^9, 10^. This study provides evidence for the mechanism by which oscillating microbubbles induce endothelial gap formation that is important for transendothelial drug delivery. We demonstrate that F-actin stress fiber severing by an oscillating microbubble followed by stress fiber recoil is crucial for creating endothelial gaps between cells within 15 s upon treatment (Fig. 2d).

We identify the F-actin cytoskeleton organization within a cell as a novel regulator of endothelial gap formation upon oscillating microbubble treatment, especially the F-actin stress fibers. Our data show that F-actin stress fiber severing by ultrasound-activated microbubbles depended on whether the fiber was within the microbubble’s maximum radius during oscillation (Fig. 4d). When microbubbles oscillate, they generate stresses: normal stresses, which are perpendicular in relation to the cell, and shear stresses, which are parallel to the cell^28^. Since the normal stresses are ∼150-fold higher than shear stresses^28^, they are most relevant in the context of stress fiber severing. For the oscillating microbubbles in our study, the normal stresses and forces were calculated, resulting in generated normal forces of 230- 8,460 nN on the F-actin stress fibers (Fig. 4f). From other methods that sever F-actin stress fibers, namely by atomic force microscopy (AFM), it is known that perpendicular forces of 200-1,000 nN^29^ are required. The lowest force value generated in our study by the oscillating microbubbles is thus in line with the lowest force value reported by AFM studies for stress fiber severing. For AFM, the 5-fold difference in the required severing forces stems from differences in fiber thickness and AFM tip geometry/sharpness^29^.

For the oscillating microbubbles, the generated forces at the used fixed ultrasound frequency and acoustic pressure varied due to differences in the oscillation amplitude of the microbubbles as a result of the variability in the initial microbubble radius. A narrower range in generated normal forces is therefore expected for monodisperse microbubbles^30, 31^ (i.e., microbubbles of the same size) in comparison to the polydisperse microbubbles we used. Currently, the models that calculate the generated force from the microbubble oscillation amplitude only hold for microbubbles that do not oscillate with a shape mode. Hence, the calculated force values for the microbubbles that oscillated with a 3^rd^ order shape mode (Fig. 4f; diamonds) are questionable and a plausible explanation why these microbubbles did not induce F- actin stress fiber severing. In addition, microbubbles that oscillate with shape modes, especially with mode ≥3, transition to a sectoral oscillation. As a result, the microbubble primarily oscillates parallel to the cell, which is the recording plane of our ultra-high-speed imaging, and has a reduced oscillation amplitude perpendicular to the cell^32, 33^. The inclusion of orthogonal ultra-high-speed imaging could provide more insight into the microbubble dynamics in 3D^34^.

Our confocal microscopy studies revealed that the severed F-actin stress fibers recoiled. This recoil is in line with the observed recoil upon F-actin stress fiber severing by laser ablation^35, 36, 37^ and AFM^29^. From these studies, it is known that severing of F-actin stress fibers can induce retraction of the entire cell for single cells without neighbors^37^ and opening of cell-cell contacts for cells in a monolayer^36^. We find that F-actin stress fiber severing by oscillating microbubbles resulted in cell-cell contact opening within 15 s upon treatment in 7 out of the 13 cells (54%). In the area of the opening, some of the CellMask dye signal is still visible (Fig. 1g). This signal suggests the presence of membrane structures, which likely is retraction fibers and migrasomes that are known to remain on substrates upon cell retraction by migration or treatment with chelating agents^38^. Tunnel formation was observed in 2 out of the 13 cells (13%) upon F-actin stress fiber severing by oscillating microbubbles. This non-uniform response is likely the result of the highly complex architecture of the F-actin cytoskeleton^36^, the location of stress fibers^37^, and the number of neighboring cells^39^. F-actin stress fibers can be removed by preincubation with the ROCK inhibitor Y-27632^40, 41^, which in our study resulted in the complete disappearance of cell-cell contact opening by oscillating microbubbles (Fig. 6d). Taken together, the data suggests that microbubble-mediated cell-cell contact opening within 15 s upon treatment is driven by F-actin stress fiber severing.

Our data shows that F-actin stress fiber severing is not essential for tunnel formation as this also occurred upon F-actin disruption only (3 out of the 4 cells, 75%) (Fig. 2). We also find that the initial and maximum F-actin disrupted area are significantly larger for tunnel formation than when the F-actin recovered after disruption (Fig. 2a,b). This tunnel formation has also been observed upon cellular dewetting^15, 16^ or applied forces by AFM^42^. From these studies, it is known that transendothelial tunnels are formed upon F-actin disruption when the apical and basal cell membranes bend and reseal around actin rings^15, 16, 43, 44^. Tunnels thereby restore the cell integrity to prevent cytosolic leakage. These tunnels can sustain for up to 20 min^45^ without leakage of cytosolic material or induction of cell death^45, 46^. When we observed a tunnel had formed, we in all cases found that the cell membranes resealed (Fig 2c). In HUVECs, a mean tunnel recovery (i.e., closure) time of 8-12 min has been reported^16, 45^, which is longer than the duration of our microscopy study. This limited time is a plausible explanation why we observed no recovery of the tunnels within our study. In our Y-27632-incubated cells, tunnel formation was reduced to only once, which is in line with another study that reported reduced tunnel formation by AFM for Y-27632-incubated cells^42^. We also find that the initial F-actin disrupted area is significantly smaller in Y-27632-incubated vs non-incubated cells when the F-actin only disrupted. These two findings combined may suggest that oscillating microbubble-induced tunnel formation is dependent on the initial F-actin disrupted area.

Because we find that F-actin disruption always occurred in conjunction with sonoporation, and the calculated microbubble oscillation amplitude for sonoporation was 0.9 µm in both cells without and with Y-27632 incubation, the F-actin disruption threshold was therefore also 0.9 µm. This sonoporation threshold of 0.9 µm is identical to the previously reported value^9^, whereas the threshold value for F-actin disruption is novel. Our finding that the sonoporation threshold is identical in cells without or with Y- 27632 incubation is in line with the same required penetration force needed by AFM in cells without or with Y-27632 incubation^42^, likely owing to the unchanged membrane tension upon Y-27632 incubation^42^. Others previously reported that the oscillating microbubble-induced membrane pore area is maximally 1.7% smaller than the disrupted F-actin area^14^. We could not confirm their finding due to the internalization of the cell membrane dye in our study, a known phenomenon^47^, which reduced the fluorescence intensity of the membrane. Of note, it is not yet clear whether oscillating microbubbles first sonoporate the cell membrane and then disrupt the F-actin that is located underneath the cell membrane^44, 48^ or whether microbubbles first disrupt the F-actin which then leads to membrane pore formation. What is known is that sonoporation can also be induced when oscillating microbubbles form microjets^49^. As these jets form perpendicular to the cell^50^ and the recording plane of our ultra-high-speed imaging was parallel to the cell, we used microbubble fragmentation as the known indicator of microjet formation^51^. Because our ultra-high-speed imaging data shows that microbubble fragmentation during oscillation was predominant when the F-actin was disrupted and absent when F-actin was not disrupted in both cells without (Fig. 3c) and with Y-27632 incubation (Fig. 6f), microjet formation was likely involved in the disruption of the F-actin and sonoporation. Our ultra-high-speed images also revealed that the oscillating microbubbles that induced F-actin disruption and sonoporation expanded more than they compressed with a median *E*/*C* ratio of 3.2 for cells without (Supplementary Fig. 2c) and 1.9 for cells with Y-27632 incubation (Supplementary Fig. 7a). These *E*/*C* ratios are higher than the previously reported median ratio of 1 for oscillating microbubbles that induced sonoporation and cell-cell contact opening in HUVEC monolayers^10^, suggesting that expansion only is not needed. The reasons for the difference in the *E*/*C* ratios could be the different acoustic pressure used in our study versus their study, namely 350 kPa versus 100, 200, 250, 300, and 400 kPa and the use of non-targeted microbubbles in our study versus αVβ3-targeted microbubbles in their study that had dampened oscillation amplitudes as a result of their internalization into the HUVECs^9^.

In our study, we also observed cell-cell contact opening >15 s, which occurred similarly in the oscillating microbubble-treated (19%) and microbubble only (24%) and ultrasound only (21%) control-treated cells, even in the absence of F-actin remodeling and sonoporation. Interestingly, all these cells were partially attached to their neighboring cells before treatment. Our findings are in line with another study that reported such cell-cell contact opening in control treatments due to external conditions like phototoxicity as a result of the confocal microscopy imaging procedure or lack of CO2 in the imaging setup^10^. This suggests that cell-cell contact opening > 15 s is an artifact of the imaging procedure rather than induced by oscillating microbubbles.

F-actin stress fibers are naturally present in endothelial cells *in vivo*, albeit it that their presence differs across the vascular tree. For example, F-actin stress fibers are present in healthy human mesenteric, mammary and epigastric arteries, but not in veins, owing to the biomechanical properties of the vascular wall^52^. In addition, the same study showed that the number of F-actin stress fibers is directly proportional to the substrate rigidity on which human arterial endothelial cells (HAECs) and HUVECs were grown *in vitro*. The reported range of the number of stress fibers per cell are in line with the amount of stress fibers we observed in our HUVECs (Fig. 4a). Conditions that increase/induce stress fiber formation in endothelial cells are adaptation to flow^53, 54^, inflammation^55, 56, 57^ and angiogenesis inhibitors^58^. Because we find that oscillating microbubble-mediated F-actin stress fiber severing induced endothelial gap formation, either openings between cells or tunnels, we propose that the efficiency of transendothelial drug delivery is highest in endothelial cells with F-actin stress fibers.

## Methods

### Lifeact-GFP transduction and cell culture

All cells were cultured at 37 °C and 5% CO2 in a humidified incubator. Human embryonic kidney 293 cells (HEK293T) were cultured in 10 cm dishes (430167, Corning) in Dulbecco’s Modified Eagle medium (DMEM, Sigma-Aldrich) supplemented with 10% fetal calf serum (16140071, Gibco, Thermo Fisher Scientific), 1% Penicillin-Streptomycin (15140122, Gibco, Thermo Fisher Scientific) and 1% L-glutamine (G7513, Sigma-Aldrich). The transfection solution was prepared using Trans-IT (MIR2304, Mirus Bio) as described by the manufacturer using a total of 15 µg of pLenti-Lifeact-GFP, pHDM-tat1b, pRC/CMV-rev1B, pHDMHgpM2, and pHDM.G constructs, a kind gift from Stephan Huveneers (Amsterdam UMC, Amsterdam, the Netherlands), diluted in a weight-ratio of 13:1:1:1:2 in 1.5 mL of Opti-MEM (31985062, Gibco). This solution was added to the 10 cm dish in a drop-wise pattern when the cells had reached ∼60% confluency. After 1 d of incubation, medium was refreshed to 5 mL of the supplemented DMEM. Transfection of the HEK293T cells was confirmed by checking for fluorescence using widefield fluorescence microscopy (Olympus IX51, Olympus). At 2 and 3 d after transfection, 5 ml of the virus-containing DMEM medium was harvested and replaced by fresh supplemented DMEM. The harvested virus-containing DMEM medium was combined and centrifuged for 5 min at 1,500 rpm (Multifuge 3s, Heraeus) after which the supernatant was filtered using a 0.45 µm filter (514-0074, VWR) to remove residual producer cells and debris. The obtained virus suspension was stored at -80° C.

Pooled human umbilical vein endothelial cells (HUVECs, #C2519A, Lonza) at passage number 3 were cultured to ∼90% confluency in 6-well plates (734-2323, VWR) in EGM-2 medium (CC-3162, Lonza). Transduction solution was prepared by mixing the virus suspension with EGM-2 medium in a ratio of 2:3 and this solution was added to the HUVECs. After 24 h, transduction was confirmed using widefield fluorescence microscopy after which the transduction solution was substituted with EGM-2 medium. Forty-eight h after transduction, selection was performed to kill the non-transfected HUVECs using 2 µg/ml puromycin (P8833, Sigma-Aldrich) in EGM-2 for 3 days. The Lifeact-GFP-HUVECs (passage number 6-8) were replated on the 50 μm thick bottom membrane of an acoustically compatible CLINIcell (Mabio)^9, 59^ and used for experiments after 2 d when the cells were fully confluent.

### Experimental setup

The microbubble-cell interaction as well as the microbubble oscillation were simultaneously imaged using a custom-built system which consisted of a Nikon A1R+ confocal microscope (Nikon Instruments)^22^ coupled to an HPV-X2 ultra-high-speed camera (Shimadzu Corp., frame rate of 10 Mfps) (Fig. 1a). During experiments, the CLINIcell was placed in the 37° C water bath with the cell-containing membrane facing up towards the microscope objective. Ultrasound was applied under an incidence angle of 45° in relation to the CLINIcell using a single element focused transducer (2.25 MHz center frequency, 76.2 mm focal length, -6 dB beam width at 2 MHz of 3 mm, V305, Panametrics-NDT, Olympus). The ultrasound pressure output was calibrated using a 1-mm diameter needle hydrophone (PA2293, Precision Acoustics). The ultrasound focal point was aligned with the optical focal point before every experiment using an pulse-echo approach and a needle tip located at the optical focal plane^60^. An arbitrary waveform generator (33220A, Agilent) and a 50 dB amplifier (ENI 2100L, Electronics & Innovation) were used to generate ultrasound pulses with a frequency of 2 MHz, pulse length of 10 cycles and peak negative pressure (PNP) of 350 kPa in water, which corresponds to an *in situ* PNP of 200 kPa after attenuation correction for the CLINIcell membrane^59^.

### Microbubble preparation

Phospholipid-coated microbubbles with a C4F10 gas core were produced in house using the previously described indirect method^61, 62^. Briefly, lipid films were created by dissolving 1,2-distearoyl-sn-glycero-3-phosphocholine (DSPC, 84.8 mol%, Lipoid GmbH), polyoxyethylene-40- stearate (PEG-40 stearate, 8.2 mol%, Sigma-Aldrich), and 1,2-distearoyl-sn-glycero-3- phosphoethanolamine-N-[carboxy(polyethyleneglycol)-2000] (DSPE-PEG(2000), 7.0 mol%, Avanti Polar Lipids) in chloroform:methanol (9:1 v/v), which was then evaporated using an argon gas-flow (Linde Gas Benelux) and freeze drying (Alpha 1–2 LD plus, Mertin Christ GmbH) overnight. Microbubble production was started by rehydrating the lipid film in 5 mL PBS which was saturated with C4F10 (F2 Chemicals) and addition of 125 µL BioTracker 400 Blue Cytoplasmic Membrane Dye (SCT109, Sigma-Aldrich) to fluorescently label the microbubble coating. Next, the lipids were dissolved using a sonication bath for 10 min and using a probe-sonicator (Sonicator ultrasonic processor XL2020, HeatSystems) at power 3 for 5 min. Microbubbles were produced by probe sonication at power 10 for 1 min, under constant flow of C4F10 gas after which the vial was filled with C4F10 gas and stored at 4 °C for a maximum of 8 days. On the day of the experiment, 1.5-2 mL microbubble suspension was washed 3-4 times by centrifugation at 400*g* for 1 min, after which the concentration was determined using a Coulter Counter Multisizer 3 (50 μm aperture tube, Beckman Coulter).

### Imaging protocol

Before imaging, the HUVECs were incubated for 10 min with CellMask^TM^ Deep Red Plasma Membrane Stain (4 µg/mL final concentration in CLINIcell, C10046, Thermo Fisher Scientific) to stain the cell membrane, followed by the addition of propidium iodide (PI, 25 µg/mL final concentration in CLINIcell, P4864, Sigma-Aldrich) and 1.8 or 2.4·10^6^ microbubbles. The CLINIcell was placed in the imaging setup with the cell-containing membrane facing up and a 5 min incubation step allowed the microbubbles to float towards the cells before imaging was started. Imaging was performed using a 100× water dipping objective (CFI Plan 100XC W, 2.5 mm working distance, Nikon Instruments) with four laser channels and filter cubes: 1) BioTracker 400 Blue excited at 405 nm and detected at 450/50 nm (center wavelength/bandwidth), 2) Lifeact-GFP excited at 488 nm and detected at 525/50 nm, 3) PI excited at 561 nm and detected at 595/50 nm, 4) CellMask^TM^ Deep Red excited at 640 nm and detected at 700/75 nm. Channel 1 and 4 were excited and detected simultaneously because there is no spectral overlap between BioTracker 400 Blue and CellMask^TM^ Deep Red. Per FOV, the imaging started with acquiring three-dimensional (3D) z-stack images with 0.33 µm z-steps using an A1 Piezo Z Drive (Nikon Instruments) at 0.12 µm/pixel (256 × 256 pixels) to image the microbubble-cell morphology in 3D prior to microbubble oscillation. Next, a 4-min time-lapse was acquired in 2D at 12-bit (i.e., fluorescent intensity ranging from 0 to 4095), 0.5 µm/pixel (FOV of 256 × 256 pixels) and 1.3 frames/s to image the cellular response to an oscillating microbubble. Ultrasound was applied ∼30 s after starting the time- lapse imaging (Fig. 1c) where the start of the ultrasound insonification was defined as *t* = 0 s. Right before the ultrasound application, the light path was automatically switched from the confocal microscope to the HPV-X2 ultra-high-speed camera to record the microbubble oscillation behavior at 10 Mfps. Upon completion of the ultra-high-speed recording, the light path was automatically switched back to the confocal to monitor the cellular response for at least 3.5 min, meaning that the confocal time-lapse imaging was shortly interrupted for 2-3 frames during ultrasound application. After time- lapse imaging, another 3D z-stack image was acquired to observe the cellular effect after microbubble oscillation in 3D. Per CLINIcell, a maximum of 15 uniformly distributed FOVs were imaged with ultrasound treatment to avoid overlapping insonification as well as a maximum of five FOVs for microbubble-only controls in the absence of ultrasound with a spacing of at least 1 cm between FOVs.

One CLINIcell was used for ultrasound only control treatments, i.e., without microbubbles. All imaging was done within 200 min. The inclusion criteria for FOVs was: i) the FOV showed a confluent monolayer in which all the cells were at least partially attached to neighboring cells; ii) there was only one microbubble in the FOV; iii) the microbubble was located on a Lifeact-GFP expressing cell that was completely in the FOV; iv) this cell had a single nucleus; and v) this nucleus did not overlap with that of any neighboring cells. FOVs were excluded when two neighboring cells were both sonoporated by one oscillating microbubble.

### Cellular response analysis

The response of the cell on which the microbubble was located before ultrasound treatment, hereafter called the cell of interest, was assessed. From the 3D z-stack and time- lapse confocal microscopy before ultrasound, the cell of interest was classified as having full junctions when its edges were fully attached to its neighboring cells or as having partial junctions when its edges were partially attached to its neighboring cells. For the controls without ultrasound, the first 30 s of the time-lapse confocal microscopy were used for the classification. After ultrasound, the cellular response was evaluated for sonoporation, endothelial gap formation and F-actin remodeling. Sonoporation of the cell of interest was analyzed and classified using a previously reported semi-automated custom-built MATLAB script^9, 10^ (R2021b, The MathWorks), which is based on the mathematical model described by Fan et al.^24^. Briefly, the cell of interest was delineated and the local increase of the PI signal upon ultrasound insonification was quantified over time. When the PI uptake curve stabilized within 120 s, the cell of interest was classified as having a resealing pore, whereas when the curve did not reach 90% of its asymptotic value within 120 s, the cell of interest was classified as having a non-resealing pore^9, 10^. In addition, if the PI fluorescent signal in the nucleus reached an intensity larger than 90% of the maximum fluorescent intensity of 4,095, at which value the DNA/RNA was considered to be saturated^10^, the pore was classified as resealing when the PI signal outside the nucleus stabilized or non-resealing when the signal kept on increasing.

Endothelial gap formation was assessed based on the 3D z-stacks and 2D time-lapse confocal microscopy recordings of the CellMask and F-actin channels. Cell-cell contact opening was classified based on changes in the cell border integrity of the cell of interest with its neighboring cells. The time of cell-cell contact opening was quantified in respect to the start of the ultrasound insonification. A transendothelial tunnel was classified when no apparent CellMask fluorescent intensity remained in the perforated area (i.e., signal comparable to background noise level).

The disrupted F-actin area was quantified based on the previously described perforated cell membrane area quantification^9, 10^ using a semi-automated custom-built MATLAB script (R2021b, The MathWorks). Briefly, the initial disrupted area in the F-actin channel in the first confocal microscopy frame after ultrasound (∼1 s) was outlined with a region of interest (ROI), after which the local change in signal over time was used to quantify the disrupted area. The F-actin responses were classified as: (a) *no disruption* when no change in the F-actin was observed, (b) *disruption and cytoskeletal recovery* when the F-actin was disrupted followed by a full or continuing recovery of the disruption area within the time-lapse, (c) *disruption only* when the F-actin was disrupted and the disrupted area either plateaued or kept on increasing during the time-lapse, or (d) *stress fiber severing and recoil* when an F-actin stress fiber was split into two in the first fluorescent frame (∼1 s) after ultrasound. All manual classifications were performed by three individual users (B.M., R.K. and Y.W.).

### Microbubble oscillation analysis

The recorded microbubble oscillations were analyzed using a custom-designed MATLAB (R2021b, The MathWorks) script as previously described^63^. From the tracked microbubble radius over time, the microbubble’s initial radius (*R0*), determined from the first 6 frames without ultrasound, maximum radius (*Rmax*), and minimum radius (*Rmin*) were obtained. Microbubbles were manually classified as fragmented when the microbubble fragmented into more than one microbubble during the oscillation. To classify the predominant shape modes, each shape mode amplitude was determined based on the tracked microbubble’s contour using a modal decomposition method^64^. The asymmetry of the microbubble oscillation was quantified by calculating the amount of expansion (*E*) relative to the amount of compression (*C*) using the formula: *E*/*C* = (*Rmax* – *R0*) / (*R0* – *Rmin*). Previously, compression-only behavior was defined by an *E*/*C* < 0.5, symmetrical oscillations by 0.5 ≤ *E*/*C* ≥ 2, and expansion-only behavior as *E*/*C* > 2^65^. The sonoporation threshold was quantified using a previously described method^10^.

### F-actin stress fiber severing and microbubble force analysis

The microbubble location was determined by selecting the microbubble center in the average BioTracker 400 Blue fluorescent signal in the time- lapse confocal microscopy recording before ultrasound using a custom-built MATLAB script (R2021b, The MathWorks). The microbubble location was also quantified as the distance from the microbubble center to the closest cell edge. In each cell of interest, stress fibers were classified as straight lines in the F-actin fluorescent channel. The total number of F-actin stress fibers and the distance between the stress fibers and microbubble center before ultrasound application were quantified using the intersection of a stress fiber with a spherical coordinate system centered at the microbubble, which was segmented into radial bands with 3 µm intervals, as shown in Fig. 4b.

The normal force (*F*) generated by an oscillating microbubble on an individual stress fiber was calculated using the normal stress model^34^, which takes into account volumetric microbubble oscillations, microbubble translation and the microbubble-boundary contact area for which we used the previously reported values^66^, as defined in equation 1:

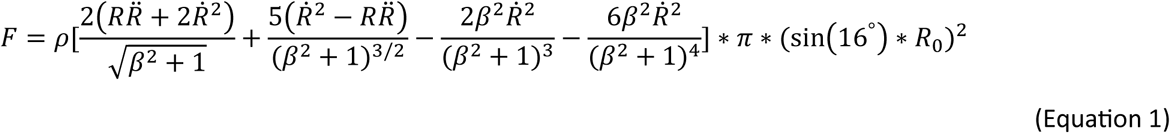

(Equation 1) where *Ṙ* and *R̈* represent the first and second-order temporal derivative of the microbubble radius *R*, p is the density of the liquid (i.e., 1·10^3^ kg/m^3^). *R0* is the resting microbubble radius, /J is the relative distance from the microbubble center to the stress fiber intended for calculation, i.e., /J = Distance⁄R_O_. The reported normal force is the maximum force (F) generated during oscillation.

### Statistics

All statistical analysis was performed in MATLAB. To assess whether data was parametric or non-parametric, the Shapiro-Wilk test of normality was used. For non-parametric data, a two-tailed Mann-Whitney U test was used with an alpha of 0.05. Parametric data was tested for equal or unequal variances using the Levene’s test followed by a two-tailed two-sample t-test with an alpha of 0.05.

Statistically significant differences are indicated in the graphs with asterisks by using * for *p* < 0.05, ** for *p* < 0.01, *** for *p* < 0.001 and **** for *p* < 0.0001. Boxplots display the median with the boxes indicating the 25th and 75th percentiles and the whiskers ranging from the minimum to maximum values excluding outliers.

### Data availability

The datasets generated in the current study are available from the corresponding authors on request.

### Code availability

Data analysis software is available from the corresponding authors on request.

## Acknowledgements

We are very grateful to Prof Dr Gijsje H. Koenderink (Department of Bionanoscience, Delft University of Technology, the Netherlands), Prof Dr Antonius F.W. van der Steen and Dr Gonzalo Collado Lara (Biomedical Engineering, Department of Cardiology, Cardiovascular Institute, Erasmus MC, the Netherlands) and Prof Dr Deborah Leckband (Chemical and Biomolecular Engineering, University of Illinois, United States of America) for insightful scientific discussions. We thank Esther van der Kamp, Dr. Ihsane Chrifi (both from Experimental Cardiology, Department of Cardiology, Cardiovascular Institute, Erasmus MC, the Netherlands), Robert Beurskens (Biomedical Engineering, Department of Cardiology, Cardiovascular Institute, Erasmus MC, the Netherlands) and Michiel Manten (Department of Experimental Medical Instrumentation, Erasmus MC, the Netherlands) for their technical assistance. This study was supported by the Applied and Engineering Sciences (TTW) (Vidi-project 17543), part of NWO, awarded to K.K.

## Author contributions

B.M., S.H. and K.K. conceived and designed the study and experiments. B.M., H.R.v.d.K., Y.W. and H.L. performed, collected data and analyzed the experiments. B.M. and K.K. wrote the manuscript. All authors discussed the results, interpreted data, reviewed and edited the manuscript.

