## Supplementary Information for "Ultrasound-activated microbubbles mediate F-actin disruptions and endothelial gap formation during sonoporation"

*Title*

**1 Biomedical Engineering, Dept. of Cardiology, Cardiovascular Institute, Erasmus MC, Rotterdam, the Netherlands**

**2 Dept. Medical Biochemistry, Amsterdam Cardiovascular Sciences, Amsterdam UMC, University of Amsterdam, Amsterdam, the Netherlands**

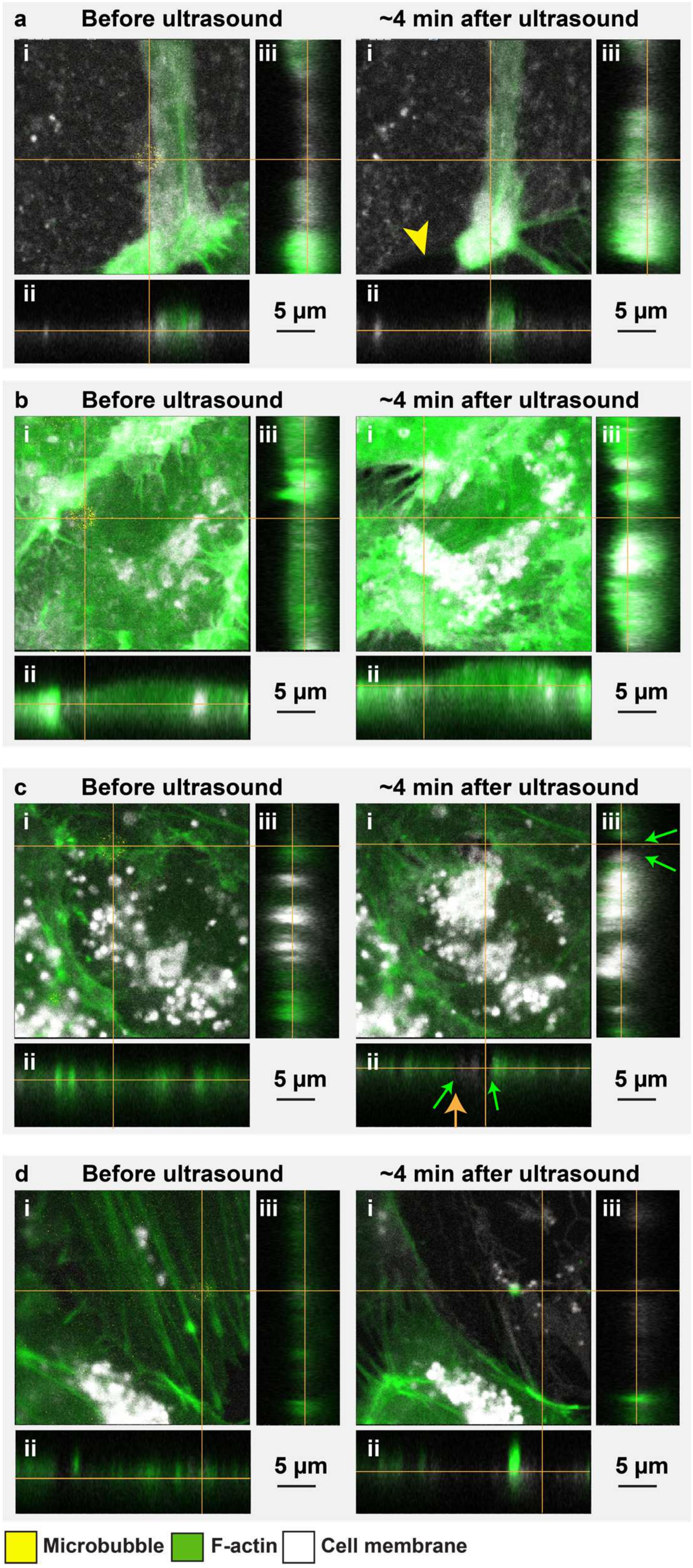


**Supplementary Figure 1 | Orthogonal views of the confocal microscopy z-stacks of the four representative F-actin remodeling examples showing the 3D morphology of the F-actin and endothelial cell membrane.** (a) No F-actin remodeling and no sonoporation. Yellow arrowhead indicates location of cell-cell contact opening. (**b**) F-actin disruption followed by cytoskeletal recovery and sonoporation. (**c**) F-actin disruption only that remained present, sonoporation and formation of transendothelial tunnel. The green arrows in the orthogonal views indicate the borders of the formed transendothelial tunnel and the orange arrow indicates location of the microbubble center before ultrasound. (**d**) F-actin stress fiber severing and recoil and sonoporation. (**a-d**) Side views have the objective imaging from (ii) the bottom or (iii) the right. Orange lines in the (i) top views indicate the cross-section of the orthogonal planes of the (ii) and (iii) side views. The location of the microbubble center is at the intersection of the orange lines, except in **c** after ultrasound where the vertical orange line is located at the formed tunnel. The (i) top views before ultrasound contain the three laser channels for the microbubble (pseudo-colored in yellow), F-actin (pseudo-colored in green), and CellMask (pseudo-colored in white), whereas the (ii) and (iii) side views and (i) top views after ultrasound contain two laser channels for F-actin and CellMask. The scale bars represent 5 µm.

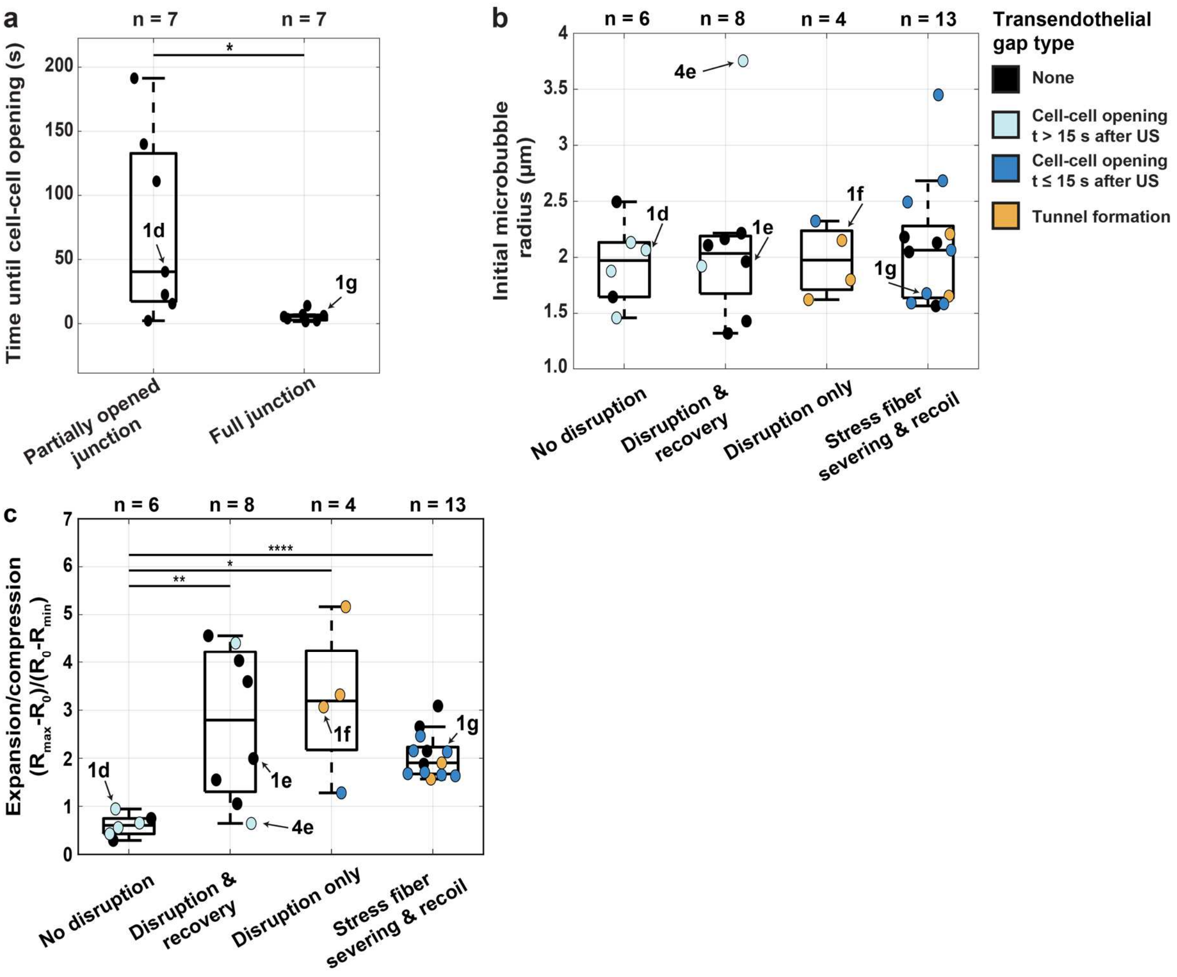


**Supplementary Figure 2 | Time till induced cell-cell contact opening and microbubble's initial radius and expansion over compression.** (**a**) Quantification of the time until cell-cell contact opening after ultrasound application in cells that were already partially attached to their neighboring cells (i.e., partially opened cell-cell junctions) before ultrasound application in comparison to cells that were fully attached to their neighboring cells (i.e., having full junctions). (**b**) Initial microbubble radius before ultrasound application. The colors of the data points correspond to the transendothelial gap types induced by the oscillating microbubbles upon ultrasound application as shown in Fig. 1d-g. (**c**) Quantification of the ratio of expansion over compression (*E*/*C*) to indicate the asymmetry of the microbubble oscillation per type of F-actin remodeling. (**a-c**) Significance is indicated with * (P < 0.05), ** (P<0.01), and **** (P<0.0001). N-numbers represent individually investigated FOVs. Boxplots represent the median and the boxes indicate the 25th and 75th percentiles with whiskers ranging from minimum to maximum values excluding outliers. Datapoints which correspond to the examples shown in Fig. 1d-g and 4e are indicated with arrows.

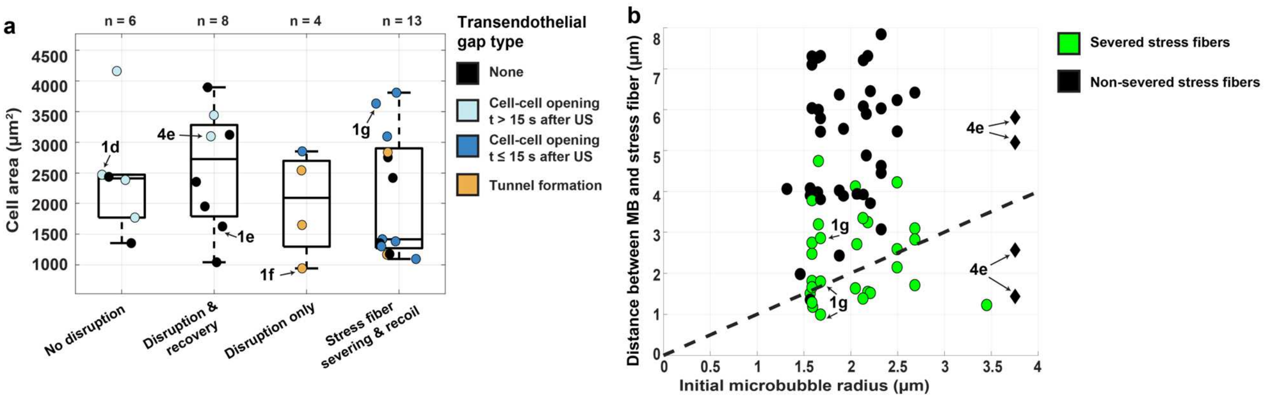


**Supplementary Figure 3 | Cell characteristics and microbubble-F-actin stress fiber morphology. (a**) Quantification of the cell area before ultrasound application. The colors of the data points correspond to the transendothelial gap types induced by the oscillating microbubbles upon ultrasound application as shown in Fig. 2d. No significant differences were observed. Boxplots represent the median and the boxes indicate the 25th and 75th percentiles with whiskers ranging from minimum to maximum values excluding outliers. (**b**) Relation between distance from the microbubble center (MB) to the surrounding F-actin stress fibers compared to the initial microbubble radius ($R_0$). Since both parameters are quantified from the center of the MB, a theoretical line (black dotted) is drawn that separates the fibers that were within reach of the initial MB location (below the line) from the fibers outside the reach of the MB location (above the line). **(a-b)** Datapoints which correspond to the examples from Fig. 1d-g and 4e are indicated with arrows.

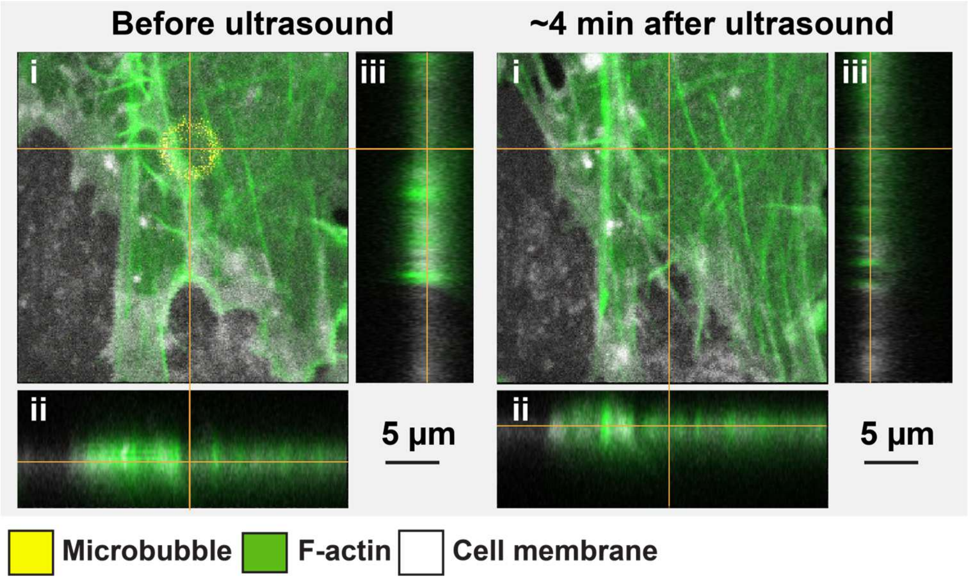


**Supplementary Figure 4. | Orthogonal views of representative example of confocal microscopy z-stacks of a microbubble with $3^{rd}$ order shape mode during oscillation showing the 3D morphology of the F-actin and endothelial cell membrane.** Side views have the objective imaging from (ii) the bottom or (iii) the right. Orange lines in the (i) top view indicate the cross-section of the orthogonal planes of the (ii) and (iii) side views. The location of the microbubble center is at the intersection of the orange lines. The (i) top view before ultrasound contains the three laser channels for the microbubble (pseudo-colored in yellow), F-actin (pseudo-colored in green) and CellMask (pseudo-colored in white), whereas the (ii) and (iii) side views and (i) top view after ultrasound contain two laser channels for F-actin and CellMask. The scale bars represent 5 µm.

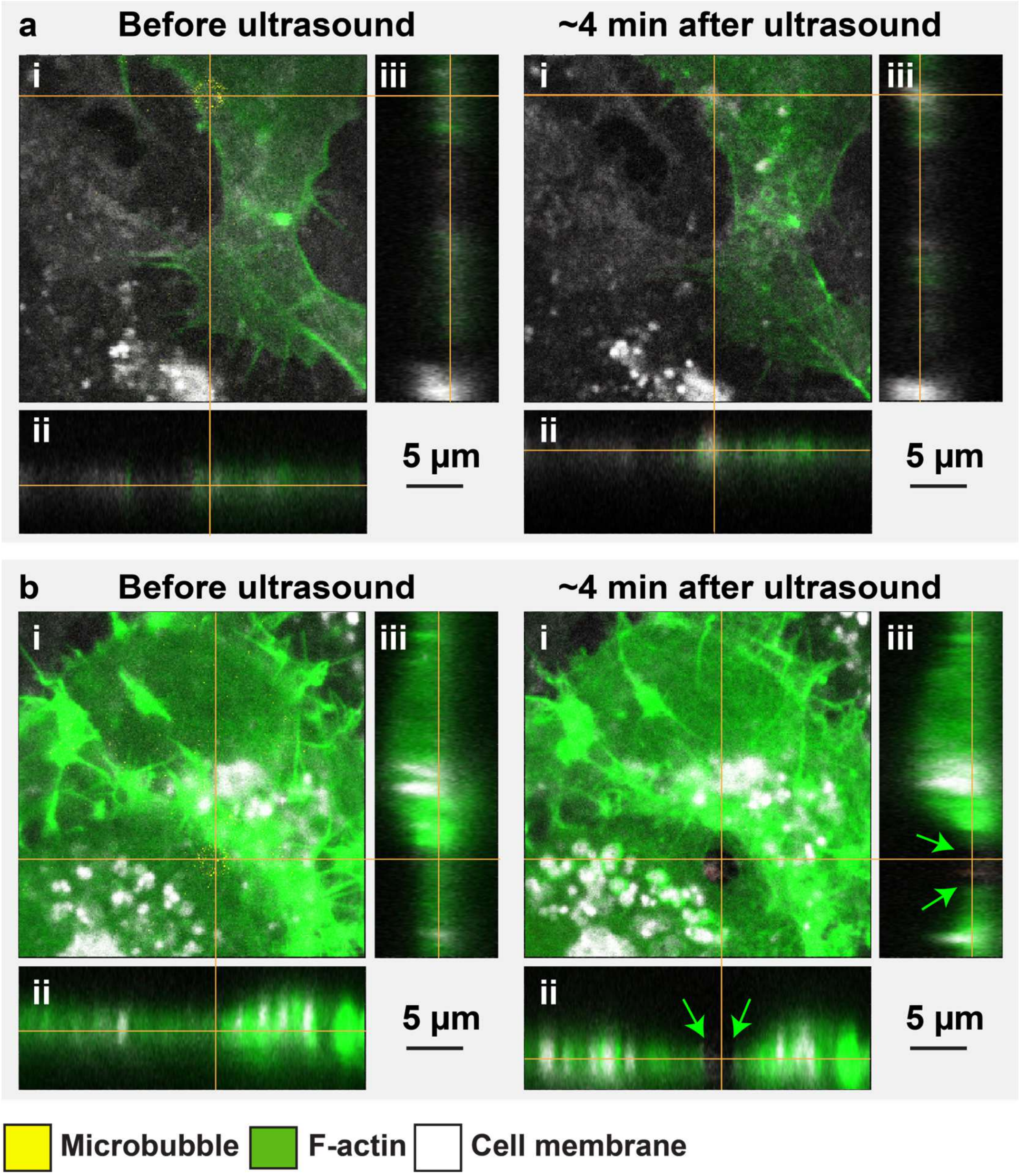


**Supplementary Figure 5 | Orthogonal views of confocal microscopy z-stacks of the two F-actin remodeling examples in the Y -27632-incubated cells showing the 3D morphology of the F-actin and endothelial cell membrane.** (**a**) F-actin disruption followed by cytoskeletal recovery and sonoporation. (**b**) F-actin disruption only that remained present, sonoporation and formation of a transendothelial tunnel. The green arrows in the orthogonal views indicate the borders of the formed transendothelial tunnel. (**a-b**) Side views have the objective imaging from (ii) the bottom or (iii) the right. Orange lines in the (i) top views indicate the cross-section of the orthogonal planes of the (ii) and (iii) side views. The location of the microbubble center is at the intersection of the orange lines. The (i) top views before ultrasound contain the three laser channels for the microbubble (pseudo-colored in yellow), F-actin (pseudo-colored in green) and CellMask (pseudo-colored in white), whereas the (ii) and (iii) side views and (i) top views after ultrasound contain two laser channels for F-actin and CellMask. The scale bars represent 5 μm.

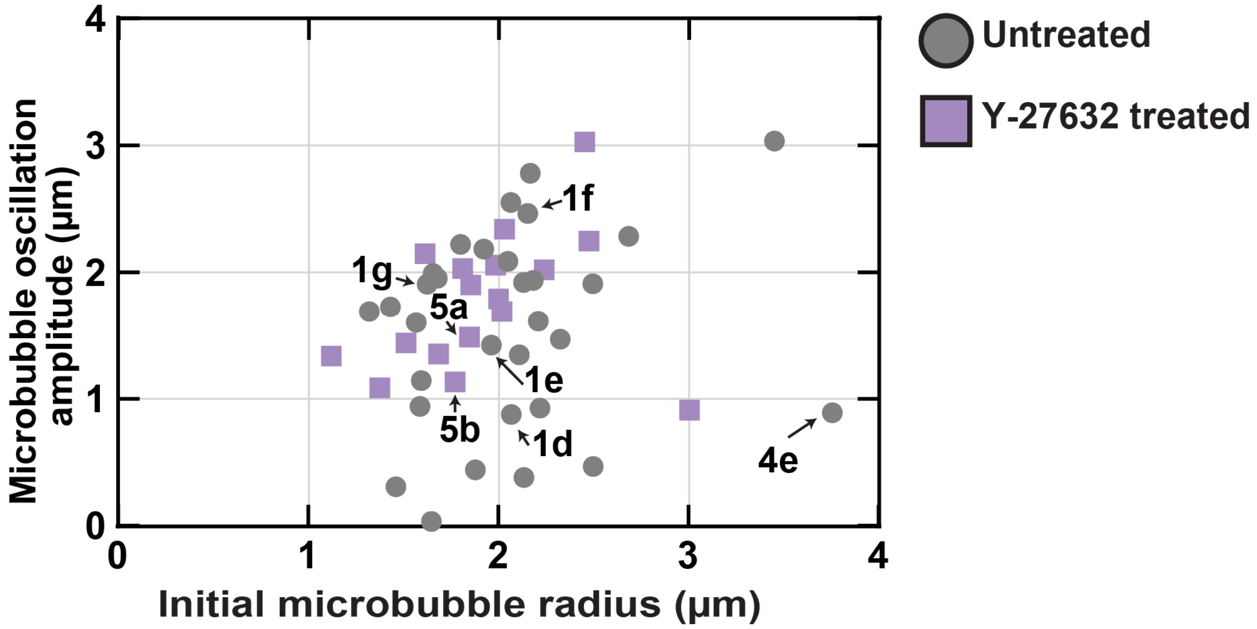


**Supplementary Figure 6 | Microbubble oscillation amplitude in cells without and with Y-27632 incubation.** The scatter dot shapes and color codes correspond to the cell preincubation type. Datapoints that correspond to the examples from Fig. 1d-g, 4e and 5a-b are indicated with arrows.

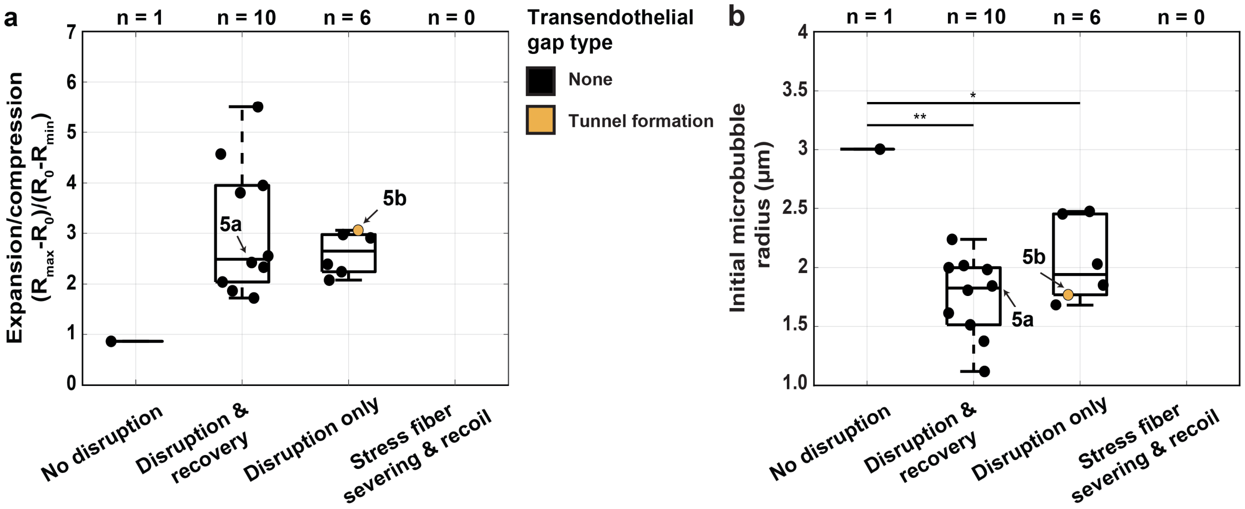


**Supplementary Figure 7 | Initial microbubble radius and microbubble expansion over compression for Y-27632-incubated cells.** (**a**) Quantification of the ratio of expansion over compression (*E*/*C*) to indicate the asymmetry of the microbubble oscillation per type of F-actin remodeling. (**b**) Initial microbubble radius before ultrasound application. (**a,b**) The colors of the data points correspond to the transendothelial gap types induced by the oscillating microbubbles upon ultrasound application. Significance is indicated with * ($P < 0.05$) and ** ($P < 0.01$). N-numbers represent individually investigated FOVs. Boxplots represent the median and the boxes indicate the 25th and 75th percentiles with whiskers ranging from minimum to maximum values excluding outliers. Datapoints which correspond to the examples from Fig. 5a-b are indicated with arrows.

**Supplementary movie 1 | Cellular response of the representative example of no F-actin remodeling and no sonoporation (corresponding to Fig. 1d).** Confocal time-lapse recording played 20× faster than real time at 21 fps where the F-actin network was visualized by lentiviral expression of LifeAct-GFP (pseudo-colored in green) and cell membrane by CellMask Deep Red (pseudo-colored in white). The microbubble location is indicated with a yellow circle. The locations where the cell was partially attached to its neighbors is indicated with asterisks. Time is indicated from the start of the video and ultrasound was applied at the 32 s timestamp, defined as t = 0 in the data analysis, during which the confocal microscopy was temporarily intercepted for HPV-X2 ultra-high-speed imaging to record the microbubble oscillation. Scale bar represents 10 µm.

**Supplementary movie 2 | Cellular response of the representative example of F-actin disruption followed by cytoskeletal recovery and sonoporation (corresponding to Fig. 1e).** Confocal time-lapse recording played 20× faster than real time at 21 fps where the F-actin network was visualized by lentiviral expression of LifeAct-GFP (pseudo-colored in green), cell membrane by CellMask Deep Red (pseudo-colored in white) and sonoporation by the uptake of the model drug propidium iodide (pseudo-colored in red). The microbubble location is indicated with a yellow circle. Time is indicated from the start of the video and ultrasound was applied at the 35 s timestamp, defined as t = 0 in the data analysis, during which the confocal microscopy was temporarily intercepted for HPV-X2 ultra-high-speed imaging to record the microbubble oscillation. Scale bar represents 10 µm.

**Supplementary movie 3 | Cellular response of the representative example of F-actin disruption only that remained present, sonoporation and formation of a transendothelial tunnel (corresponding to Fig. 1f).** Confocal time-lapse recording played 20× faster than real time at 21 fps where the F-actin network was visualized by lentiviral expression of LifeAct-GFP (pseudo-colored in green), cell membrane by CellMask Deep Red (pseudo-colored in white), and sonoporation by the uptake of the model drug propidium iodide (pseudo-colored in red). The microbubble location is indicated with a yellow circle. Time is indicated from the start of the video and ultrasound was applied at the 38 s timestamp, which is defined as t = 0 in the data analysis, during which the confocal microscopy was temporarily intercepted for HPV-X2 ultra-high-speed imaging to record the microbubble oscillation. Scale bar represents 10 µm.

**Supplementary movie 4 | Cellular response of the representative example of F-actin stress fiber severing and recoil, sonoporation and cell-cell contact opening <15 s (corresponding to Fig. 1g).** Confocal time-lapse recording played 20× faster than real time at 21 fps where the F-actin network was visualized by lentiviral expression of LifeAct-GFP (pseudo-colored in green), cell membrane by CellMask Deep Red (pseudo-colored in white) and sonoporation by the uptake of the model drug propidium iodide (pseudo-colored in red). The microbubble location is indicated with a yellow circle. Time is indicated from the start of the video and ultrasound was applied at the 24 s timestamp, which is defined as t = 0 in the data analysis, during which the confocal microscopy was temporarily intercepted for HPV-X2 ultra-high-speed imaging to record the microbubble oscillation. Scale bar represents 10 µm.

**Supplementary movie 5 | Cellular response of the representative example of F-actin disruption and cell-cell contact opening <15 s induced by a microbubble located in a preexisting gap between neighboring cells**. Confocal time-lapse recording played 20× faster than real time at 21 fps where the F-actin network was visualized by lentiviral expression of LifeAct-GFP (pseudo-colored in green), cell membrane by CellMask Deep Red (pseudo-colored in white) and sonoporation by the uptake of the model drug propidium iodide (pseudo-colored in red). The microbubble location is indicated with a yellow circle. Time is indicated from the start of the video and ultrasound was applied at the 26 s timestamp, which is defined as t = 0 in the data analysis, during which the confocal microscopy was temporarily intercepted for HPV-X2 ultra-high-speed imaging to record the microbubble oscillation. Scale bar represents 10 µm.

**Supplementary movie 6 | Ultra-high-speed recording of microbubble oscillation and fragmentation (corresponding to Fig. 1g and 3d).** Time is indicated from the start of the ultra-high-speed recording (played at 21 fps) and ultrasound (2 MHz: 350 kPa PNP, 10 cycles) was applied at 6,500 ns. Scale bar represents 2 µm.

**Supplementary movie 7 | Ultra-high-speed recording of microbubble oscillating with 3rd order shape mode (corresponding to Fig. 3e).** Time is indicated from the start of the ultra-high-speed recording (played at 21 fps) and ultrasound (2 MHz: 350 kPa PNP, 10 cycles) was applied at 6,500 ns. Scale bar represents 2 µm.

**Supplementary movie 8 | Cellular response of the example of F-actin disruption in between two stress fibers followed by recovery and sonoporation (corresponding to Fig. 4e).** Confocal time-lapse recording played 20× faster than real time at 21 fps where the F-actin network was visualized by lentiviral expression of LifeAct-GFP (pseudo-colored in green), cell membrane by CellMask Deep Red (pseudo-colored in white) and sonoporation by the uptake of the model drug propidium iodide (pseudo-colored in red). The microbubble location is indicated with a yellow circle. Partial junctions are indicated with asterisks. Time is indicated from the start of the video and ultrasound was applied at the 44 s timestamp, which is defined as t = 0 in the data analysis, during which the confocal microscopy was temporarily intercepted for HPV-X2 ultra-high-speed imaging to record the microbubble oscillation. Scale bar represents 10 µm.

**Supplementary movie 9 | Cellular response of the representative example of F-actin disruption and recovery and sonoporation in a Y-27632-incubated cell (corresponding to Fig. 5a).** Confocal time-lapse recording played 20× faster than real time at 21 fps where the F-actin network was visualized by lentiviral expression of LifeAct-GFP (pseudo-colored in green), cell membrane by CellMask Deep Red (pseudo-colored in white) and sonoporation by the uptake of the model drug propidium iodide (pseudo-colored in red). The microbubble location is indicated with a yellow circle. Time is indicated from the start of the video and ultrasound was applied at the 32 s timestamp, which is defined as t = 0 in the data analysis, during which the confocal microscopy was temporarily intercepted for HPV-X2 ultra-high-speed imaging to record the microbubble oscillation. Scale bar represents 10 µm.

**Supplementary movie 10 | Cellular response of the example of F-actin disruption only that remained present and sonoporation in a Y-27632-incubated cell (corresponding to Fig. 5b).** Confocal time-lapse recording played 20× faster than real time at 21 fps where the F-actin network was visualized by lentiviral expression of LifeAct-GFP (pseudo-colored in green), cell membrane by CellMask Deep Red (pseudo-colored in white) and sonoporation by the uptake of the model drug propidium iodide (pseudo-colored in red). The microbubble location is indicated with a yellow- circle. Time is indicated from the start of the video and ultrasound was applied at the 24 s timestamp, which is defined as t = 0 in the data analysis, during which the confocal microscopy was temporarily intercepted for HPV-X2 ultra-high-speed imaging to record the microbubble oscillation. Scale bar represents 10 µm.
